# Unsupervised Voxel-based Segmentation reveals a Landscape of Bacterial Ribosome Large Subunit Early Assembly

**DOI:** 10.1101/2022.11.09.515851

**Authors:** Kai Sheng, Ning Li, Jessica N. Rabuck-Gibbons, Xiyu Dong, Dmitry Lyumkis, James R. Williamson

## Abstract

Ribosome biogenesis is a complex but efficient process in rapidly growing bacteria, where assemble a functional 70S ribosome takes ~ 2 min, involving participation of 3 rRNAs, 50 r-proteins and dozens of assembly factors. In vitro reconstitution using different subsets of large subunit (50S, LSU) proteins with rRNAs, pioneered by Nierhaus lab, resulted in the Nierhaus assembly map, embodying the cooperativity and dependency for binding of LSU r-proteins to 23S rRNA. Critically absent from the Nierhaus map is the underlying folding of the rRNA that creates the binding sites for the r-proteins. In addition, the relationship of the observed cooperativity in vitro to the co-transcriptional assembly in cells remains to be determined.

Pre-50S intermediates accumulate at low temperature in ΔdeaD, a DEAD-box helicase implicated in 50S assembly. We solved 21 pre-50S density maps from intermediate-containing fractions using cryo-EM. In the newly solved maps, we discovered the earliest intermediate ever reported, consisting of domain I at the 5’-end of 23S rRNA. To probe the mechanism behind the maps during assembly, we developed a novel density map segmentation and dependency analysis method. Ten cooperative assembly blocks were identified from segmentation of the maps, and these were organized into a block dependency map. This is the first time the dependencies on folding of rRNA helices and ribosomal protein binding could be integrated into a complete assembly map. In addition, we showed how the exit tunnel is folded on the solvent side, serving as a scaffold for 50S maturation. Using this new segmentation analysis method, we revisited previously reported bL17-depletion and ΔsrmB datasets. Most remarkably, the other two datasets are also consistent with the block dependency, implying a unified early assembly pathway and flexible maturation landscape in early 50S biogenesis.

## Introduction

Ribosome biogenesis is a complex but efficient process in rapidly growing bacteria, where assembly of a functional 70S ribosome takes ~ 2 min[1, 2], involving participation of 3 rRNAs, 50 r-proteins and dozens of assembly factors [3–6]. *In vitro* reconstitution using various subsets of large subunit (50S, LSU) proteins with rRNAs, resulted in the Nierhaus assembly map[7–9], embodying the cooperativity and dependency for binding of LSU r-proteins to 23S rRNA. Critically absent from the Nierhaus map is the underlying folding of the rRNA that creates the binding sites for the r-proteins. In addition, the relationship of the observed cooperativity *in vitro* to the co-transcriptional assembly in cells remains to be determined.

The structure of the complete 50S subunit provides few clues to the assembly pathway. The 23S rRNA secondary structure is organized into 6 domains based on phylogenetic analysis of secondary structure, but these domains are highly interdigitated in the complete subunit. Using a genetic depletion of bL17[10], a series of 13 intermediates was identified using cryo-EM for the later stages of assembly. Analysis of this set of particles revealed 5 assembly blocks, not specifically organized by secondary structure domains, with evidence for both parallel and sequential folding of blocks of RNA. The earliest intermediate in that study consisted of approximately the solvent side half of the 50S subunit. Genetic manipulation of assembly factors, like Srmb, ObgE, RbgA, *etc*. [11–13], has generated additional structural data that reveals additional details of the mechanism of late-stage assembly of the intersubunit half of the 50S subunit.

The deletion strain ΔdeaD, has a severe growth defect at low temperature, and the sucrose gradient profile for this strain shows significant accumulation of a pre50S peak. Cryo-EM analysis of the pre-50S peak from the DdeaD strain identified the earliest intermediate yet identified, consisting of domain I and three associated r-proteins. The analysis of this data was facilitated by an improved workflow for heterogeneous reconstruction and iterative subclassification using template-free ab-initio model building in CryoSPARC. Further, we also developed a new toolbox of unsupervised feature extraction and electron density segmentation to identify assembly blocks, based on single voxel behavior across a set of maps, which enables the subsequent cooperativity and dependency analysis.

Overall, we generated a set of 21 pre50S density maps from the ΔdeaD dataset, and we applied our segmentation and dependency analysis method to identify 10 cooperative assembly blocks. The set of blocks was organized into a block dependency map, that demonstrated for the first time the integrated interdependency of organization of rRNA helices and protein binding. The process by which the exit tunnel is formed was revealed during assembly of the solvent half of the subunit, which then serves as a scaffold for 50S maturation. With the folding blocks of the entire subunit in hand, we revisited the previously reported bL17-depletion and ΔsrmB datasets[10, 12].Most remarkably, the other two datasets are also consistent with the block dependency derived from the deaD dataset, which implies a unified early assembly pathway and a malleable maturation landscape in early 50S biogenesis.

## Results

### Iterative ab initio subclassification reveals new LSU assembly intermediates

DeaD is a cold shock protein in *E. coli*, with several annotated functions involving mRNA stability and association with the 50S ribosomal subunit[14–16]. The ΔdeaD strain grown at 19 C resulted in the accumulation of pre50S particles in the sucrose gradient profile. The whole cell and pre50S fraction proteomic data showed that a lack of of DeaD at low temperature results in a defect of ribosome assembly without alteration of r-protein expression (SI).

The pre-50S fractions from the ΔdeaD strain were subjected to single particle cryoEM data collection using a similar approach to the iterative subclassification strategy previously reported[17]. Our previous approach was implemented using Frealign, but we developed a robust alternative implementation for iterative ab-initio subclassification using CryoSPARC, as described in the methods. Analysis using this protocol resulted in 21 distinct particle maps for the ΔdeaD dataset, shown in Figure 1a.

**Fig. 1.**
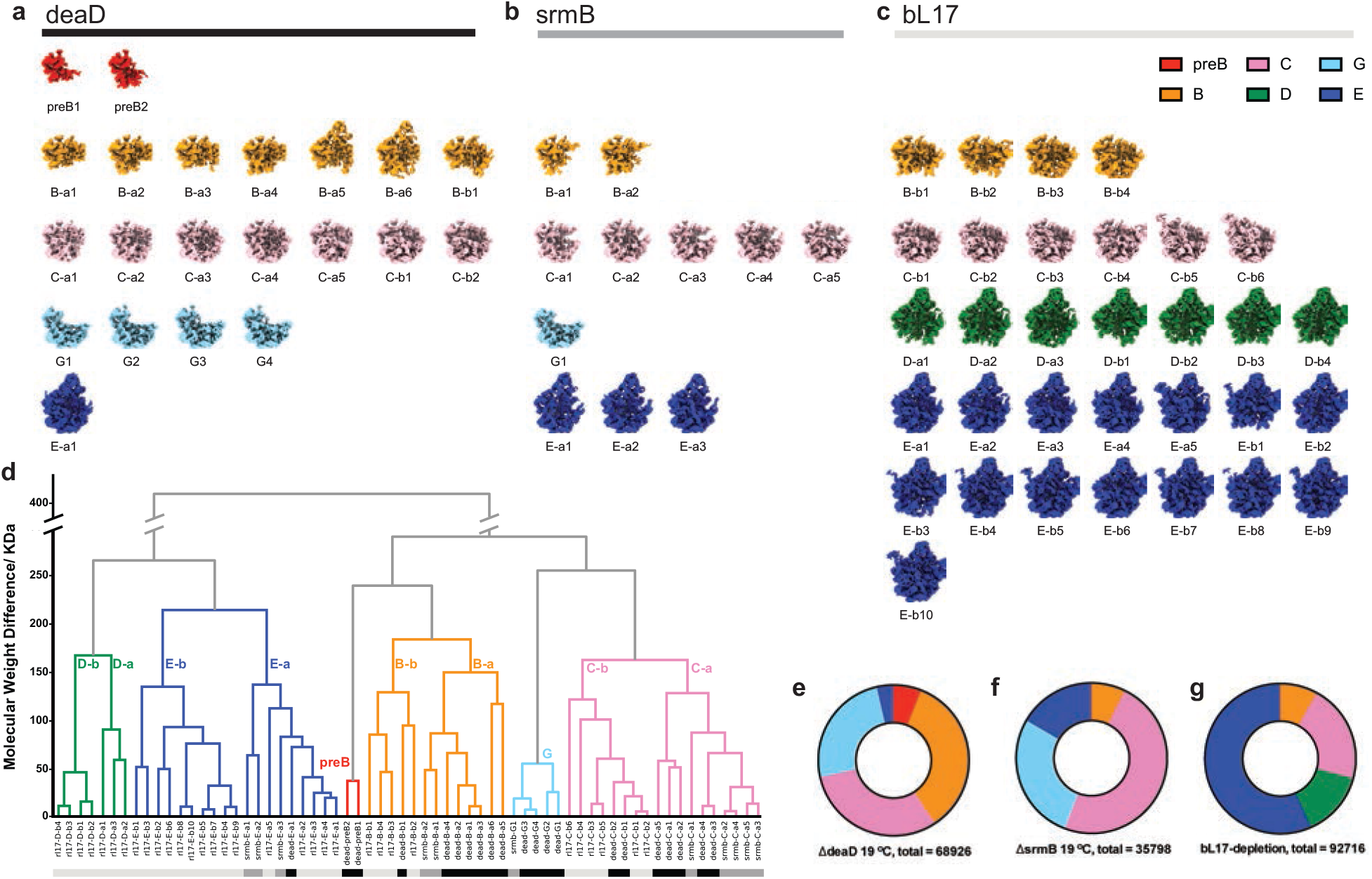
Assembly intermediate density maps from three datasets. Density maps reconstructed from (**a**) ΔdeaD, (**b**) ΔsrmB and (**c**) bL17-depletion datasets, colored according to classes obtained from hierarchical analysis in (**d**) The Euclidean distance matrix, based on molecular weight in kDa, was calculated among density maps, and the dendrogram resulting from hierarchical clustering is displayed, with the 6 main class branches colored accordingly. (**e-g**) Particle distribution among the main classes for the three datasets.

To compare with prior results, we re-analyzed two previously reported datasets using this updated protocol, one with intermediates from depletion of bL17 [10], and one with intermediates from a ΔsrmB strain [12], resulting in 32 and 11 distinct particle maps, respectively, shown in Figure 1b,c. The combined set of 64 maps were compared to identify similar classes of particles among the three individual sets of maps. The Euclidean distance matrix was calculated for the total set of 64 maps, thresholded at 1% of the maximum voxel intensity. Agglomerative hierarchical clustering was performed using this matrix, which grouped the maps into 6 major classes (Figure 1d), with the class distributions for the three datasets shown in Figure 1e,f,g. Some of the new maps align well to the B, C, and E classes previously observed in the bL17 depletion strain [10, 17], while the D-class is only observed in the bL17 dataset. Two new classes observed in the ΔdeaD dataset were labeled as pre-B and G.

The newly discovered pre-B-1 and pre-B-2 classes, exclusively found in the ΔdeaD dataset, represent the earliest intermediates yet observed, corresponding roughly to domain I or domain I+III. There is a new class related to the previous C class that is primarily observed in the ΔdeaD dataset (25% of the particles in Figure 1e, metadata in Table S2), which we assign to the new G class that lacks a large portion of domain II helices, as well as uL13, bL20 and bL21. The ΔdeaD particles mainly consist of the early B, C and G classes, while the E class only represents a small fraction of the particles. The ΔdeaD dataset exhibited the largest breadth of early intermediates, ranging from the smallest intermediate yet observed, corresponding to ~600 nucleotides of domain I, to a mature E class, which connects well to previous data focusing on the late assembly process.

### Unsupervised voxel-based segmentation of density maps from ΔdeaD intermediates reveals ten early assembly blocks

We developed a novel procedure to segment the maps using the dimensional reduction tools PCA[18] and UMAP[19], in combination with the HDBSCAN[20, 21] algorithm for cluster identification (see Methods). The PCA-UMAP-HDBSCAN segmentation method identified ten early assembly blocks as a basis set for the experimental maps(Figure 2). Briefly, the set of 21 maps from ΔdeaD were aligned and resampled to the same grid, thresholded at 1% of their maximum signal, and the resulting 21 × 114,392 matrix of voxels with intensity above threshold in any dataset was used to generate 21 principal components. Plotting the first two principal components, PC1 and PC2, showed distinguishable features, but the clusters are not readily separated (SI). Some features can be extracted by thresholding a given PC at 1s, such as the positive elements of PC2 as the base region or the negative elements of PC3 as domain I. (Extended data Fig.1 ab). In general, the desired contiguous density segments would be linear combinations of the PCs, but there is no straightforward method to solve for those segments. In addition, features in higher PCs are noisy and hard to identify, and voxels in these PCs cannot be unambiguously assigned to a single structural feature (Extended data Fig.2). Direct application of UMAP on the voxel intensities suffered from similar shortcomings in identifying contiguous segments (SI). In contrast, subjecting the PCA reconstructed data to dimensionality reduction using UMAP gave rise to readily interpretable groups of voxels, as shown in Figure 2a.

**Fig. 2.**
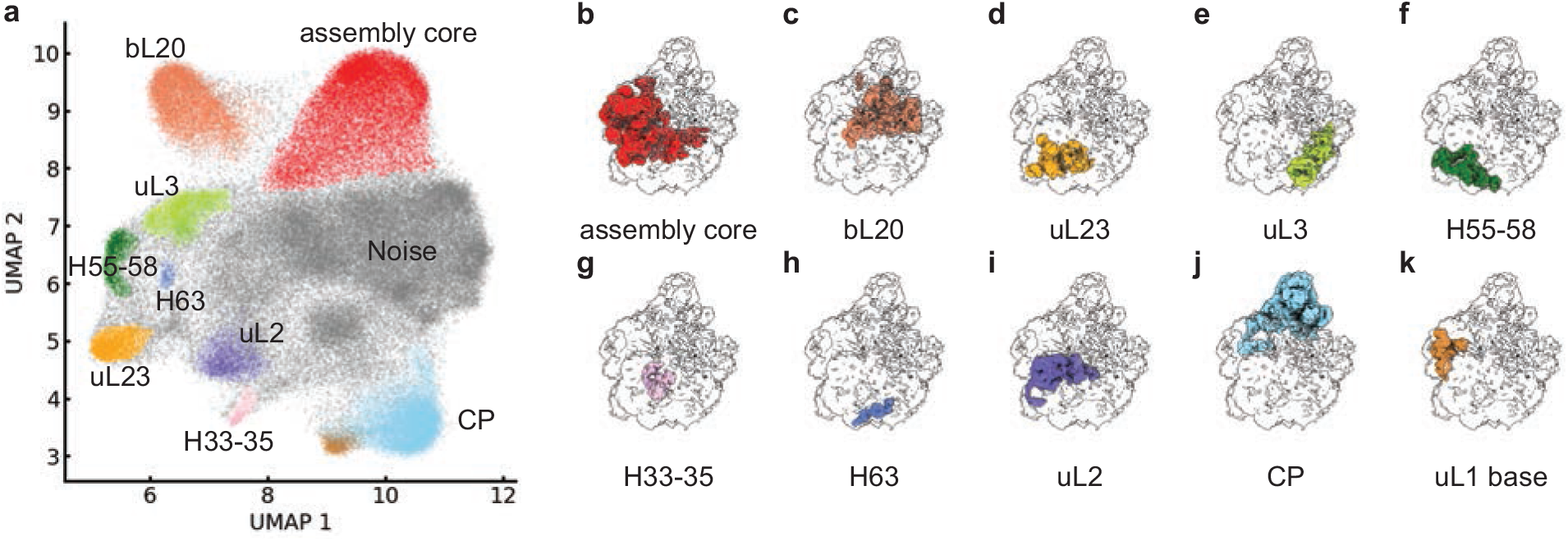
Assembly blocks derived from segmentation using PCA-UMAP-HDBSCAN. **a**) Voxels above a 1% threshold are well organized in UMAP space, with 10 contiguous volume blocks extracted by clustering with HDBSCAN colored (see Methods/SI). **b-k)** Projections of the clusters into 3D space overlaid on the 50S subunit 1% threshold mask (black outline), colored according to **a)**.

Finally, HDBSCAN was used to cleanly and quickly resolve and identify clusters of voxels in the UMAP representation in an unsupervised manner (Figure 2a). The resulting clusters do in fact correspond to contiguous regions of electron density that serve as a basis set for the 21 maps from the ΔdeaD dataset, as shown in Figure 2b-k. The sequential application of PCA-UMAP-HDBSCAN analysis provides the cleanest most intuitive segmentation of the set of density maps, representing a powerful and general template for analyzing sets of maps from heterogeneous cryo-EM reconstruction.

The set of blocks was subjected to occupancy analysis using the RNA helices and r-proteins, allowing assignment of the blocks to particular structural elements (Extended data Fig.3). Nearly all of the RNA and protein elements are uniquely assigned to a single block, confirming that the blocks can be used as a convenient basis set for assembly. In order to understand the relationship among the newly defined blocks and to compare distinct features in individual maps, we performed a new occupancy analysis of the 21 intermediate maps based on the 10 basis blocks, resulting in a 10×21 matrix shown in Figure 3a. The blocks names were based on prominent RNA or protein features, Figures 2ab, 3a.

**Fig. 3.**
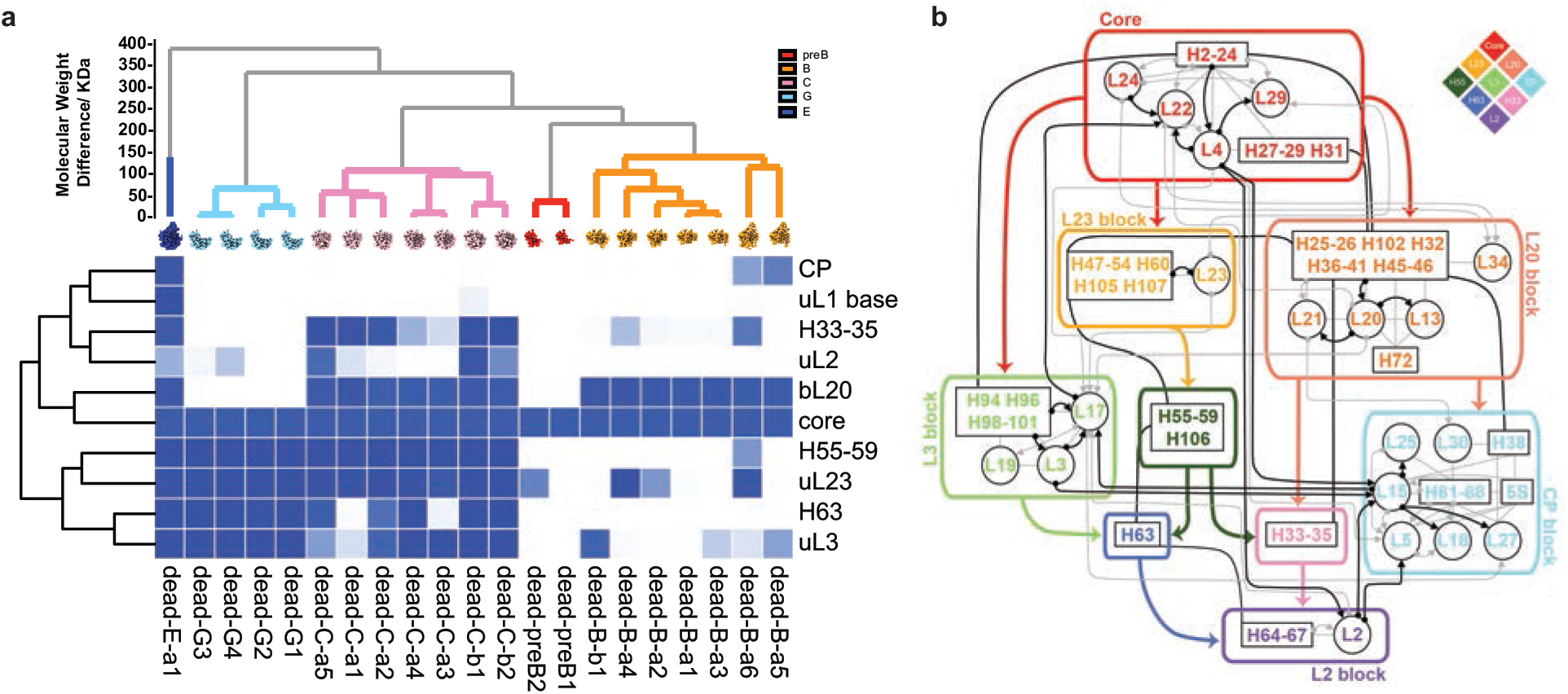
Occupancy matrix of assembly blocks with the resulting block dependency map. **a**) Electron density occupancy of 21 intermediate density maps from ΔdeaD in terms of the 10 assembly blocks, used for dependency analysis. **b**) Block dependencies were determined using a quadrant analysis of the occupancy matrix in **a)** (see SI). All of the 23S and 5S rRNA helices are outlined in black boxes, with connections between elements in primary structure in solid black lines. Black/grey arrows show dependencies from the Nierhaus map as strong/weak interaction. The major block dependencies inferred from **a)** are shown as bold colored arrows. The diamond schematic diagram of the blocks, used in Figure 4, is shown as an inset at the upper right.

### Assembly Block dependency maps cooperativity for early 50S assembly

A folding block is operationally defined as a set of voxels intensities that are correlated across the 21 maps, and are thus considered to be a cooperative assembly unit. The composition of the blocks is schematically shown in Figure 3a, superimposed on the protein binding dependencies from the original Nierhaus map. The dependency among the blocks is evaluated by first making a scatterplot for each pair of rows of the occupancy matrix, then inferring the dependency using a quadrant analysis (Extended data Fig.4 & Methods). The uL1 base block was omitted from the dependency analysis, as it only occurred once in the ΔdeaD dataset. After pruning the redundant edges in the dependency graph (SI), the most direct dependencies between blocks are retained, and they are shown in colored bold arrows in Figure 3b.

The assembly Core is a prerequisite for all subsequent assembly blocks, and in particular is the sole precursor to consolidation of the bL20 or uL23 blocks. These blocks constitute domain I, II and III, respectively, providing strong support for the natural cotranscriptional direction of assembly. The uL3 block represents a significant portion of domain VI, and can be organized by either the core + bL20 (in early B classes) or the core + L23 (in G classes). We do not have evidence of direct folding of the uL3 on the assembly core, likely due to the rapid folding kinetics of the L20 and L23 blocks. The association of domains I, II, III, and VI forms the majority of the solvent half of the 50S subunit, and this group serves as the template for final assembly of the CP, the stalks, and ultimately the peptidyl transferase center. There are three small blocks consisting entirely of RNA, the H55-59/106 block which is part of domain III, the H63 block which is part of domain IV, and the H33-35 block, which is part of domain II. Finally, there is a large block corresponding to the CP, and the L2 block which is part of domain IV. (Extended data Fig.5, detailed block descriptions in SI). The remaining parts of the 50S subunit not covered by the 10 blocks correspond to the last folding steps forming the active site, that has been informed by previous work on the bL17 depletion strain. Moreover, the newly identified assembly blocks subdivided previously reported assembly, which illuminated key features of early 50S assembly (Extended data Fig.6).

### Placing RNA helices in the early stages of the assembly map

The comprehensive assembly map shown in Figure 3b, includes both the r-proteins and the rRNA helices defined in the secondary structure. The well-known Nierhaus assembly map established the basis for thermodynamic cooperativity among the LSU proteins binding to the 23S rRNA [7–9]. For the first time, it is possible to intertwine RNA secondary structure elements with the LSU proteins to produce an RNA-protein assembly map. This is particularly revealing for the earliest stages of assembly, and it is now clear that the assembly primarily proceeds in the 5’-3’ direction, consistent with a co-transcriptional organization of the folding blocks[22].

### The minimal requirement for CP formation

The dependency graph in Figure 3b shows that the minimal requirement for CP formation is the assembly core docked with the bL20 block. The deaD-B-a5 particle is the smallest intermediate with CP formed, composed of the assembly core, the bL20 block and a partially formed uL3 block. In dead-B-a6, it is larger than dead-B-a5 while the uL3 block is not formed at all. With this evidence, which is consistent with the dependency graph, we conclude that the CP formation only requires the assembly core and bL20 block, which is continuous primary sequence from domain I to domain II except for H38cp and H42-44 (base for L7/12 stalk), after the H38bd is formed, the H38cp will recruit corresponding proteins, 5S and part of domain V (H81-88) and form an intact CP(Extended data Fig.7d).

### Formation of exit tunnel in early intermediates

The discovery of early intermediates reveals the layer-by-layer formation of exit tunnel from the solvent side towards the inter-subunit side, which finally forms the peptidyl transfer center (PTC). We split the exit tunnel (ET) into two parts: ET_solvent_ and ET_PTC_ (Extended data Fig.8a). It is more relevant to discuss the ET_PTC_ formation in the bL17 depletion dataset since they contain more mature structures, while the less mature set of ΔdeaD intermediates, allows a focus on the ET_solvent_ formation. By analyzing the structure of the secM and vemP peptide trapped on a translating ribosome [23, 24], we generated a list of 50S contacts in each of the assembly blocks within 5 A of the trapped peptide (Table S3).

From the assembly blocks, we can easily assign the resolved residues in ET_solvent_ into the assembly core, bL20, uL23 block and H33-35 block. Intermediates that do not contain all three of these blocks, the pre-B, G and B (except dead-B-a6) class and part of the early C class, do not have a fully structured ET_solvent_. In the dependency graph, we know formation of H33-35 is dependent on bL20 block and H55-58, that implies H55-58 is also a prerequisite for ET_solvent_ formation. For example, though the matured base region formed in the G class intermediates, none have the bL20 block formed, and so they never have the H33-35 block formed, resulting in incomplete ET_solvent_ formation. All of the C classes have the H55-59, uL23 block and bL20 blocks, and they only need the H33-35 formation to complete the ET_solvent_ during the maturation.

These results differ significantly with the Steinberg analysis of the evolution of the 23S rRNA from the primordial PTC [25], in which domain I (assembly core), domain II (bL20 block) and domain III (uL23 block) form after PTC formation, in the evolutionary sense. Presumably, the folding of the proto-ribosome was organized around the PTC, and there was likely an important stage in evolution, after insertion of the domains into the proto-PTC, where the assembly process was reorganized from forming the PTC first, to forming the ET_solvent_ first as a scaffold on which to build the PTC.

### Different perturbations reshape the ribosome assembly landscape

Given the observed dependencies in Figure 3b, a set of 29 possible combinations of allowable structures can be enumerated, and of these, 14 were observed in the ΔdeaD dataset. These structures can be organized into a putative assembly mechanism, connecting similar structures according to the dependencies, as shown in Figure 4a. We also calculated the block occupancy for the other two datasets (Extended data Fig.9). With the occupancy matrix, we validate the quadrant analysis across the three datasets, using ΔdeaD thresholding criteria (Extended data Fig.10). The quadrant analysis perfectly matched the ΔdeaD results. Across the ΔdeaD, ΔsrmB, and bL17 datasets, 21 of the 29 possible structures are observed, and there are no combinations that violate the dependencies based on the ΔdeaD data alone (Figure 3b). Comparing different datasets on the pathway, the ΔdeaD intermediates are distributed earlier than the other three, and there are few intermediates containing CP blocks.

**Fig. 4.**
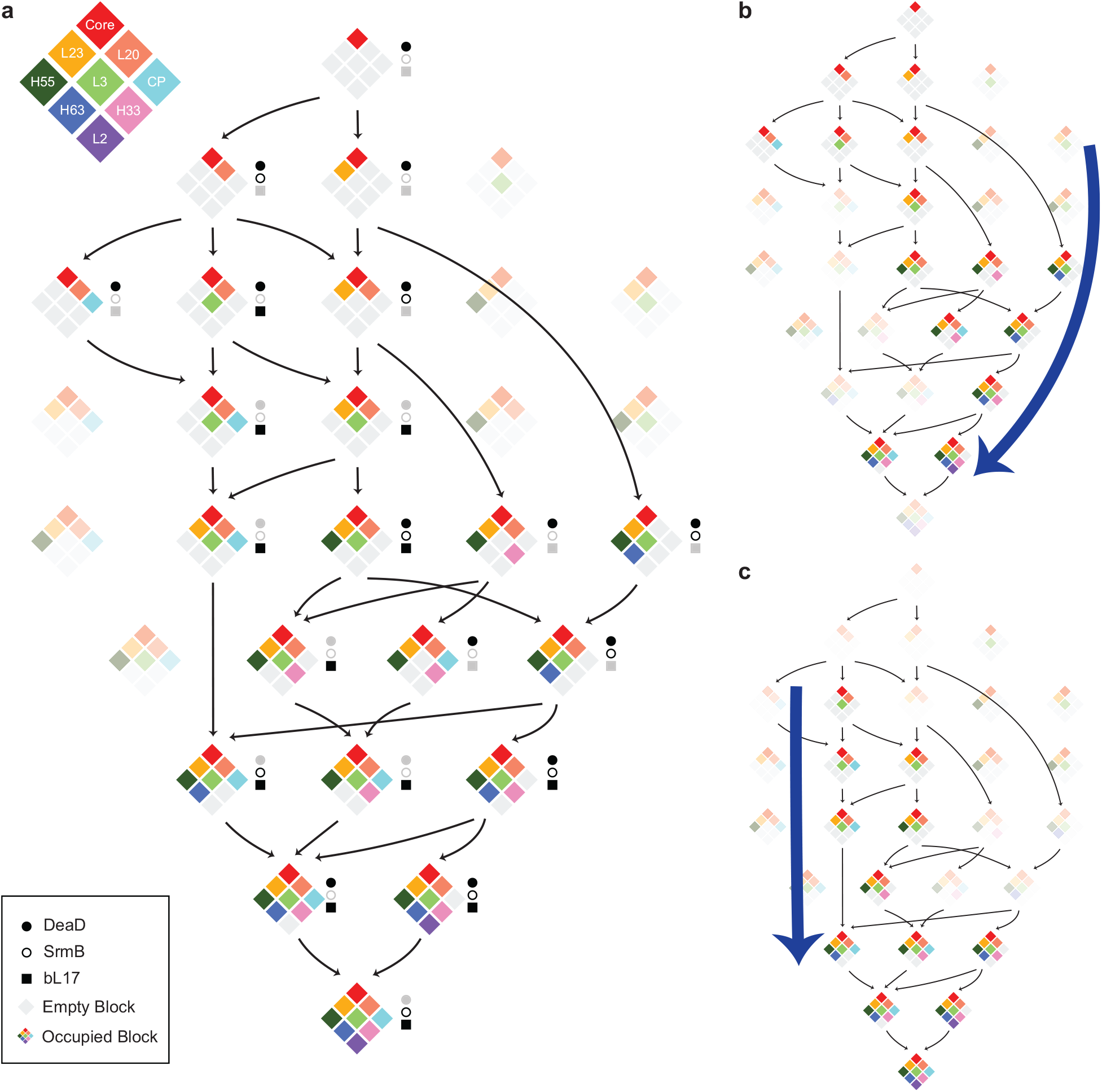
Assembly pathways using the diamond block schematic. **a**) Based on the block dependencies in Figure 3b, 29 possible intermediates are arranged in a layout based on increasing of block number from top to bottom. Intermediates observed in the datasets are shown in color with any unoccupied blocks shown in gray. The presence of an intermediate in ΔdeaD, ΔsrmB or bL17-depletion datasets is shown by black closed circles, open circles and squares, respectively. There are 8 combinations consistent with the block dependencies that are not observed, indicated by faded intensities. Arrows connect the nearest precursors, with disassembly not allowed, requiring the assembly core as parent node. Display of intermediate found in ΔdeaD (**b**) and bL17-depletion (**c**) datasets. Blue bold arrows highlight the change in flux through same set of intermediates in the different datasets. (See SI for the ΔsrmB intermediate pathway, which is similar to ΔdeaD)

The other two datasets populate intermediates not observed in the ΔdeaD dataset, *** but consistent with the DdeaD dependencies, which implies a universal block-wise parallel pathway for the assembly mechanism (Figure 4a). Interestingly, the ΔdeaD and ΔsrmB datasets share a similar assembly path, through the unique G class compared to the bL17-depletion strain. We’ve shown that ΔsrmB intermediates have depletion of uL13 resulting in a defect in the bL20 block formation. Also, the bL20 block is essential for formation of the CP block, so in the ΔsrmB dataset, there are fewer intermediate with the CP block formed at the very beginning, flowing through the right side of the landscape in Figure 4b (ΔsrmB pathway in Extended data Fig.11). In contrast, bL17 is at the bottom of the mature 50S and is thought to play an important role in the blocks of the base region (uL2 block, uL3 block, H33-35 and H63 block), so the bL17-depletion intermediates have defects in the base blocks’ formation, the CP forms earlier intermediates accumulate with an incomplete base region, and mostly go through the left part of the assembly landscape in Figure 4c. Although there no knowledge of the substrate for deaD during ribosome assembly, the ΔdeaD strain shares many intermediates with the ΔsrmB strain, which implies the two helicases operate on substrates that are present at similar times during assembly.

### Continuous learning of 50S folding dependency with new datasets

We have shown that the essential features of the early stage 50S assembly pathways for datasets from different perturbations can be represented using segments derived from the ΔdeaD dataset alone. As the field progresses, additional information will become available through novel perturbations, and it becomes interesting consider how the analysis differs when additional datasets are included. Performing the PCA-UMAP-HDBSCAN analysis on the three combined datasets with 64 intermediate density maps revealed a set 18 assembly blocks (namely, blk01 to blk18). To compare to the 10ΔdeaD blocks, occupancy analysis was performed for the new set of blocks (Extended data Fig. 12), revealing 8 blocks that were perfectly aligned in common. (Extended data Fig. 13a-d, f-h). Interestingly, a small segment was separated from the bL20 block from ΔdeaD that corresponded to uL13 and H25 (Extended data Fig. 13e). The depletion of uL13 was a feature of intermediates in the ΔsrmB strain, implying that the site of action of SrmB may be related to related to H25. In a similar manner, bL17 was segmented out from uL3 block owing to including bL17-depletion dataset (Extended data Fig. 13i). Further, new blocks were identified in the 3-dataset analysis, such as H67-69 and uthe L10/uL11 stalks (Extended data Fig. 14cd) that are important in late assembly stages, and are not present in the early ΔdeaD datasets. The blk14 even includes a non-native density for assembly factor yjgA (Extended data Fig. 14g), which is also feature of a subset of bL17-depletion intermediates, capturing the distinct dumbbell shape of YjgA surrounded by bL31 and H74/80/93 as a docking site for YjgA. The PCA-UMAP-HDBSCAN approach can robustly identify distinct and common features, for both native and nonnative density, in a complex assembly landscape, and provides for continuous learning of the ribosome biogenesis pathway as more observations are included using different perturbations.

By analysis of the particles that accumulate in the ΔdeaD strain at low temperature, we identified a diverse set of intermediates that span the entire pathway for assembly of the 50S subunit. These include a series of intermediates that include the earliest particles composed roughly of domain I at the 5’ end of the subunit, proceeding with assembly of the solvent portion of the peptide exit tunnel, prior to assembly of the intersubunit face and PTC. We developed a novel segmentation method for a set of density maps that allows segment identification, and used those segments to develop an assembly pathway for the entire subunit. This segmentation can be readily applied to other datasets resulting from different perturbations, providing a unifying set of intermediates across datasets with differing fluxes through the pathway. Overall, these data and the subsequent analysis provide a comprehensive view of the overall assembly of the 50S subunit that integrates a significant body of data from decades of research into a coherent assembly map containing all r-proteins and RNA helical elements.

## Methods

### Bacterial strains and plasmid construction

Strains BW25113 E.coli (WT) and BW25113(ΔdeaD) from the Keio Knockout Collection were purchased from the E.coli Genetic Stock Center[26]. The pHSL-deaD, homoserine lactone (HSL) -inducible deaD expression plasmid, was generated by Gibson cloning from pHSL-rplQ[10], replacing the coding region of rplQ with deaD coding sequence in pHSL. The ΔdeaD-pHSL-deaD strain was obtained by transformation of pHSL-deaD into strain ΔdeaD.

### Cell growth and sucrose gradient purification for ribosome particles

WT, ΔdeaD and ΔdeaD-pHSL-deaD strains were inoculated in LB medium and grown overnight, then diluted into fresh LB at 20 °C. Either 0 or 2.5 nM HSL was added into the ΔdeaD-pHSL-deaD strain during cell culture. Cells were harvested at OD600 ~ 0.4 by centrifugation at 4000 rpm for 15 min, followed by lysis in Buffer A (20 mM Tris-HCl pH 7.5, 100 mM NH4Cl, 10 mM MgCl2, 0.5 mM EDTA, 6 mM β-mercaptoethanol) and 20 U/ml DNase I (Sigma) by a mini bead beater using 0.1-mm zirconia/silica beads (3×60 seconds pulses with 1 min on ice in between). Insoluble cell debris and beads were then removed by two centrifugation steps: 16,000 rpm (31,000 g) for 10 min, transferring the supernatant to a new tube and then again 16,000 rpm for 90 min. The clarified cell lysates (10 A_260_ units) were loaded onto a 33 mL 10-40 % w/v sucrose gradient (50 mM Tris-HCl 7.8, 100 mM NH_4_Cl, 10 mM MgCl_2_, 6 mM β-mercaptoethanol) then centrifuged in a Beckman SW32 rotor at 26,000 rpm for 16hours at 4 °C. Gradients were fractionated using a Brandel gradient fractionator. Based on the UV 254 nm trace, gradient fractions corresponding to the pre-50S peak were collected and combined. To prepare the fractions for cryo-EM analysis, 3x volumes of buffer A were added prior to concentration in a 100 kDa cutoff concentrator (Amicon) 3 times to eliminate sucrose and to equilibrate to buffer A.

### Electron microscopy sample preparation and data collection

The purified pre-50S sample was diluted to 0.6 mg/ml with buffer A, and 3 uL of the sample was applied to a plasma cleaned gold grid in a humidified CP3 chamber (FEI). Grids were manually frozen in liquid ethane, and single particle data was collected on a Thermo Fisher Scientific Titan Krios electron microscope operating at 300k eV equipped with a Gatan K2 Summit detector using the Leginon software[27], with a pixel size of 1.31 Å at 22,500X magnification. A dose of 33 to 35 e^−^/ Å^2^ across 50 frames was used for a dose rate of ~5.8 e/pix/sec. To overcome problems of preferred orientation of particles on the grid, data was collected at the tilt of −20° [28]. A total of 1031 micrographs were collected.

### Electron microscopy micrograph processing

Data pre-processing was performed in the Appion pipeline[29] including individual programs that are cited. Frames were aligned using MotionCor2 software[30], and the contrast transfer function (CTF) for all micrographs was performed with CTFFind4.1[31]. A total of 322,187 particles from the dataset were picked with auto picking in RELION3. The particles were extracted with bin of 2 and fed into 2D classification to remove 30S, 70S and spurious particles, then further cleaned with 3D classification using a C-class 50S intermediate as a template[17]. After 3D classification from RELION, classes that did not produce an interpretable map were eliminated. The resulting 273,729 particles were exported and analyzed in CryoSPARC3 [32]. Particle stacks for bL17-depletion and ΔsrmB were prepared using the same procedure on previously acquired micrographs [10, 12], resulted in 123,804 and 273,620 particles respectively.

### Iterative classification with ab initio reconstruction in CryoSPARC and hierarchical analysis

For each dataset, the resulting particle stacks were imported to CryoSPARC and directly subjected to *ab inito* reconstruction, asking for 4 classes with default parameters. Each resulting interpretable class was subjected to another round of *ab initio* reconstruction using the same parameters. This procedure was performed iteratively until the particle number in a class was less than 2,000. All reconstructions with fewer than 2,000 particles were subjected to *ab initio* reconstricution, asking for 1 class before 3D refinement in CryoSPARC (See SI for ΔdeaD dataset examples). All refined density maps were aligned and resampled to a same 50S ribosome reference (bL17-depletion dataset E) in ChimeraX. All maps with resolution below 10 Å were discarded and the remaining resampled density maps were thresholded at intensity 1.00. Pairwise difference maps were calculated for the binarized (thresholded at 1.00) maps the sum of the difference map A-B and difference map B-A for hierarchical clustering using the Ward linkage.

Maps were displayed with the resulting dendrogram, and pairs of maps with a difference of < 10 KDa were merged into one class, recalculating the map using ab initio reconstruction and 3D refinement. This step is important as similar classes can emerge from hiding at various stages of the iterative subclassification. (SEE SI) Finally, a hierarchical clustering analysis was performed across all three datasets in the same way to allow ready comparison of maps from the different datasets.

### Segmentation using PCA-UMAP-HDBSCAN with ΔdeaD intermediates and three datasets

The 21 resampled DdeaD intermediate maps were thresholded at 1% of their maximum signal and, and the set of 114,392 voxels with nonzero intensity in at least one map resulted in a 21 × 114,392 intensity array. Principle component analysis (PCA in Scikit-Learn)[18, 33] was performed on this array, giving PCA transformed matrix of the same dimensions. UMAP[19] analysis was performed on the PCA matrix using 2 components with 100 nearest neighbors using the Canberra metric, resulting in a 2 × 114,392 matrix, projecting each voxel above the intensity threshold into a UMAP_1,2_ space. In the UMAP_1,2_ space, HDBSCAN (min_cluster_size = 100, min_samples = 100)[20, 21] was performed to assign voxels to individual clusters, resulting 9 blocks representing contiguous regions of density in Cartesian space, and one noise block. The 9 blocks were considered as assembly blocks, corresponding to a basis set of voxels that have correlated intensities in the input set of 21 maps. Blocks were named according to salient structural or compositional features. The CP and L1 base blocks were further separated with another round of HDBSCAN (min_cluster_size = 10, min_samples = 10). All resulting assembly blocks were cleaned by dust filtering in ChimeraX[34] prior to use in occupancy analysis. (See SI for algorithm parameters discussion). Similarly, 64 intermediates from three datasets generated a 64 × 140,545 matrix. PCA and UMAP were performed direct on the first dimension of the matrix. HDBSCAN(min_cluster_size = 200, min_samples = 10) was performed to extract 18 assembly blocks.

### Occupancy and dependency analysis

The Occupancy of each assembly block was calculated for each of the density maps thresholded at intensity 1.00. Briefly, the number of voxels above threshold are counted in each block then normalized to the total number of voxels in the block. The occupied fraction for each block is then normalized to the core block occupancy in each density map.

The dependency between any pair of blocks (*i,j*) was obtained by quadrant analysis of a scatter plot of the occupancy for block *i* on the x-axis and block *j* on the y-axis (Extended data Fig. 4a). The dashed binarization lines for the horizontal and vertical directions were calculated by the following equation (eq.1),

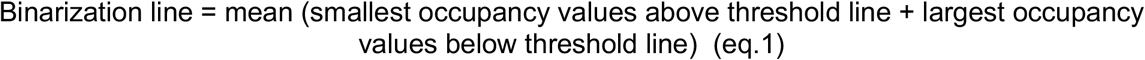

* threshold line is defined in Extended data Fig. 16.

The x and y binarization lines divide the scatter plot into 4 quadrants: QI = lower left, QII =lower right, QIII = upper left, QIV = upper right. To infer the relationship between block *i* and *j*, the number of points in each quadrant were counted for the scatter plot. The relationship between block *i* and *j* falls into one of three scenarios (Extended data Fig. 4a). With points only in QI and/or QIV, blocks *i* and *j* are correlated. With dots in both QII/QIII, blocks *i* and *j* are not correlated. With dots only in QI/QII/QIV or QII/QIV, block *j* should depend on block *j* (red scatter plots in Extended data Fig. 4b). With dots in only QI/QIII/QIV or QIII/QIV, block *i* should depend on block *j*. (blue scatter plots in Extended data Fig. 4b). If block *i* depends on block *j*, an arrow from *j* to *i* will be drawn in the dependency map. The comprehensive dependency plot is now ready for pruning with defined rules (SI) with networkx package[35].

## Code Accessibility

The code for volume curation (alignment and resample), hierarchical analysis, PCA-UMAP-HDBSCAN, occupancy analysis and quadrant-dependency analysis can be found at: https://github.com/ks277/2022_50S_landscape_paper (updating in process, scripts can be obtained by emailing the author)

## ACKNOWLEDGMENTS

This work was supported by grants from the National Institutes of Health GM136412 and GM053757 to JRW.

---

**Extended data Fig. 1.**
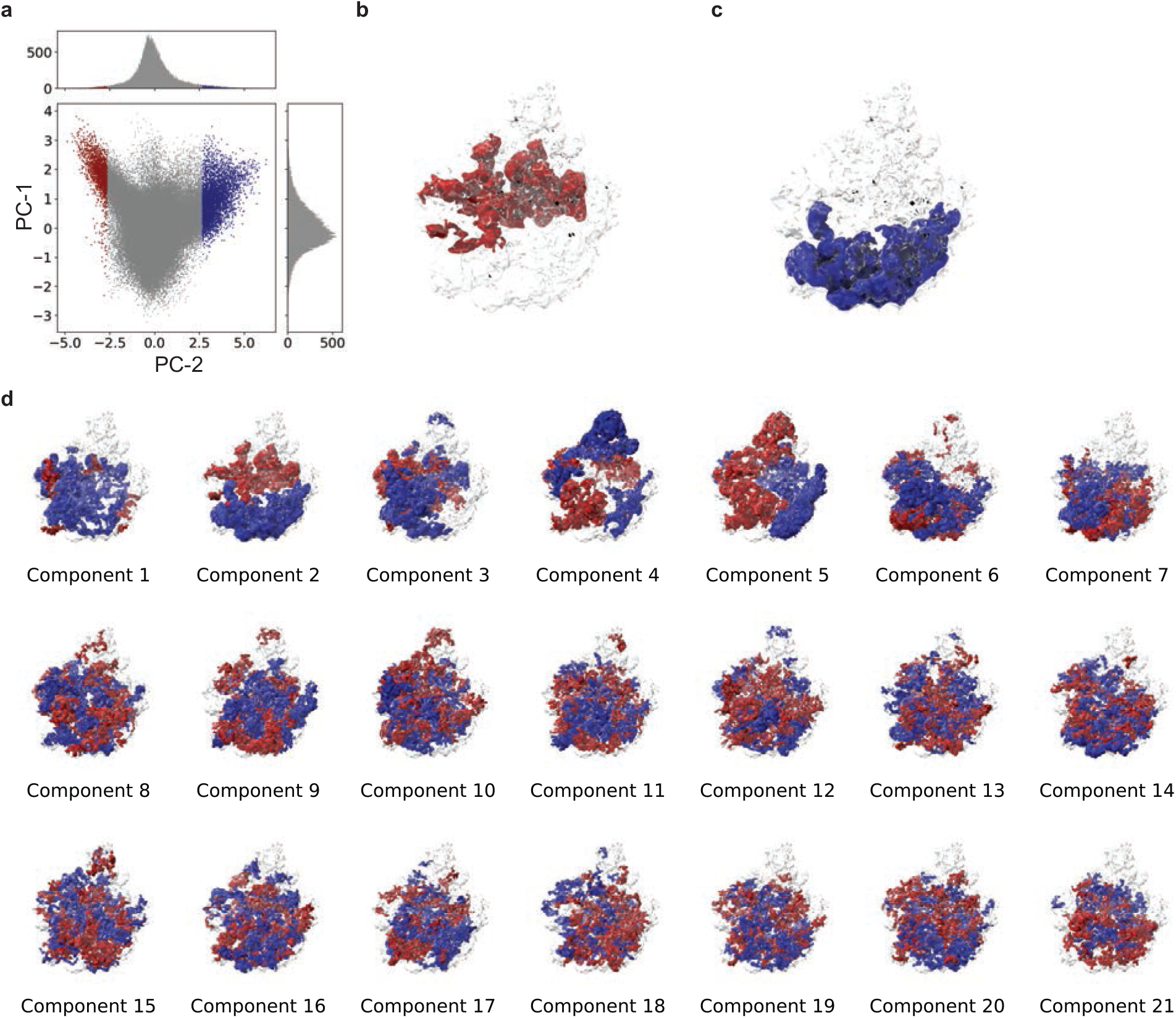
Principal Component Analysis (PCA) of voxels above threshold in the ΔdeaD dataset. **a)** Scatterplot of the transformed value of each voxel in PC-1/PC-2 coordinates, with histograms of the marginal distributions at top and right. The extraction process is illustrated with the second principal component, PC-2. Voxels were thresholded on the PC-2 axis after thresholding at +/− 1σ, resulting in volumes for a negative feature **b)** and a positive feature **c). d**) Features extracted from PC-1 to PC-21 using the same criteria in **a**). All volumes in 3D space are overlaid on the 50S subunit 1% threshold mask calculated from PDB 4ybb (transparent white).

**Extended data Fig. 2.**
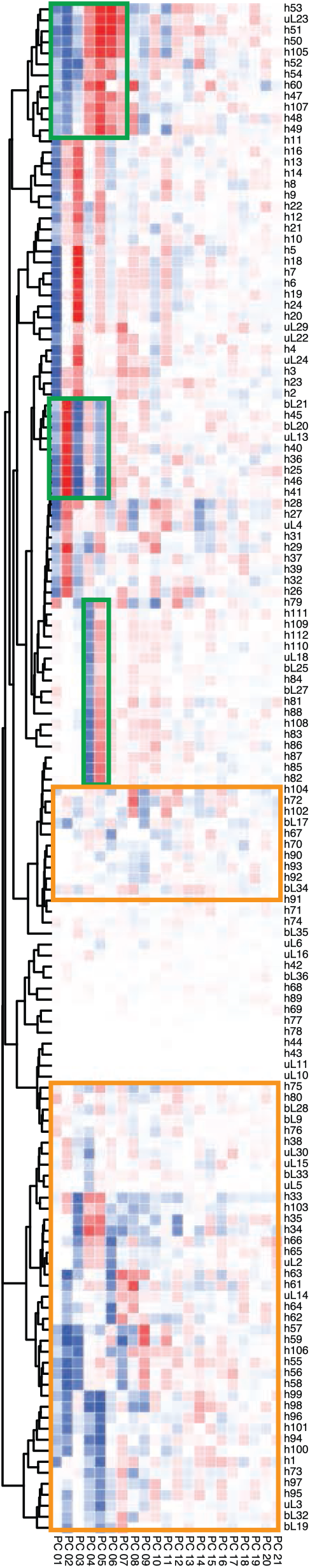
Occupancy matrix of positive and negative features extracted in Extended data Fig. 1. Voxel occupancy of PCA extracted features in 139 segments from 50S r-proteins and helices are calculated, with positive values in blue and negative values in red, and displayed in a clustered heat map. The x-axis is ordered according to the PCs, and the y-axis was ordered by hierarchical clustering. The challenges for feature extraction are highlighted by green boxes, where segments are extracted multiple times in different PCs as both negative and positive features. Orange boxes highlight cases where it is difficult to identify segments, which is particularly problematic for the higher PCs. In general, the higher PCs are primarily noise and the lower PCs are linear combinations of the desired features, but there is no straightforward way to extract the features from the PCs.

**Extended data Fig. 3.**
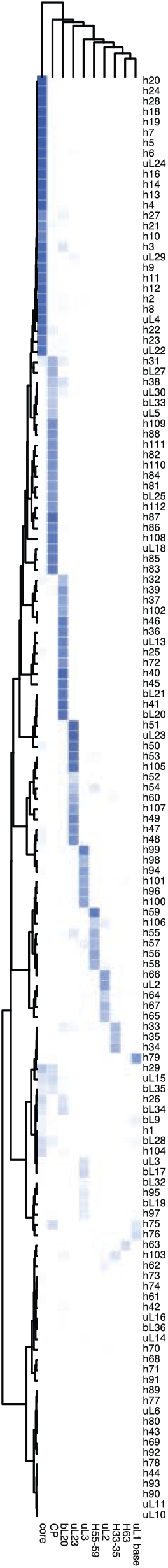
Occupancy matrix of blocks resulting from PCA-UMAP-HDBSCAN with in ΔdeaD dataset. Voxel occupancy of PCA-UMAP-HDBSCAN extracted blocks in segments from 50S r-proteins and helices are calculated and displayed in a clustered heat map. The x- and y-axes ordered by clustering, using Euclidean distance with Ward linkage. Extraction of features from the PCA-UMAP-HDBSCAN representation is straightforward.

**Extended data Fig. 4.**
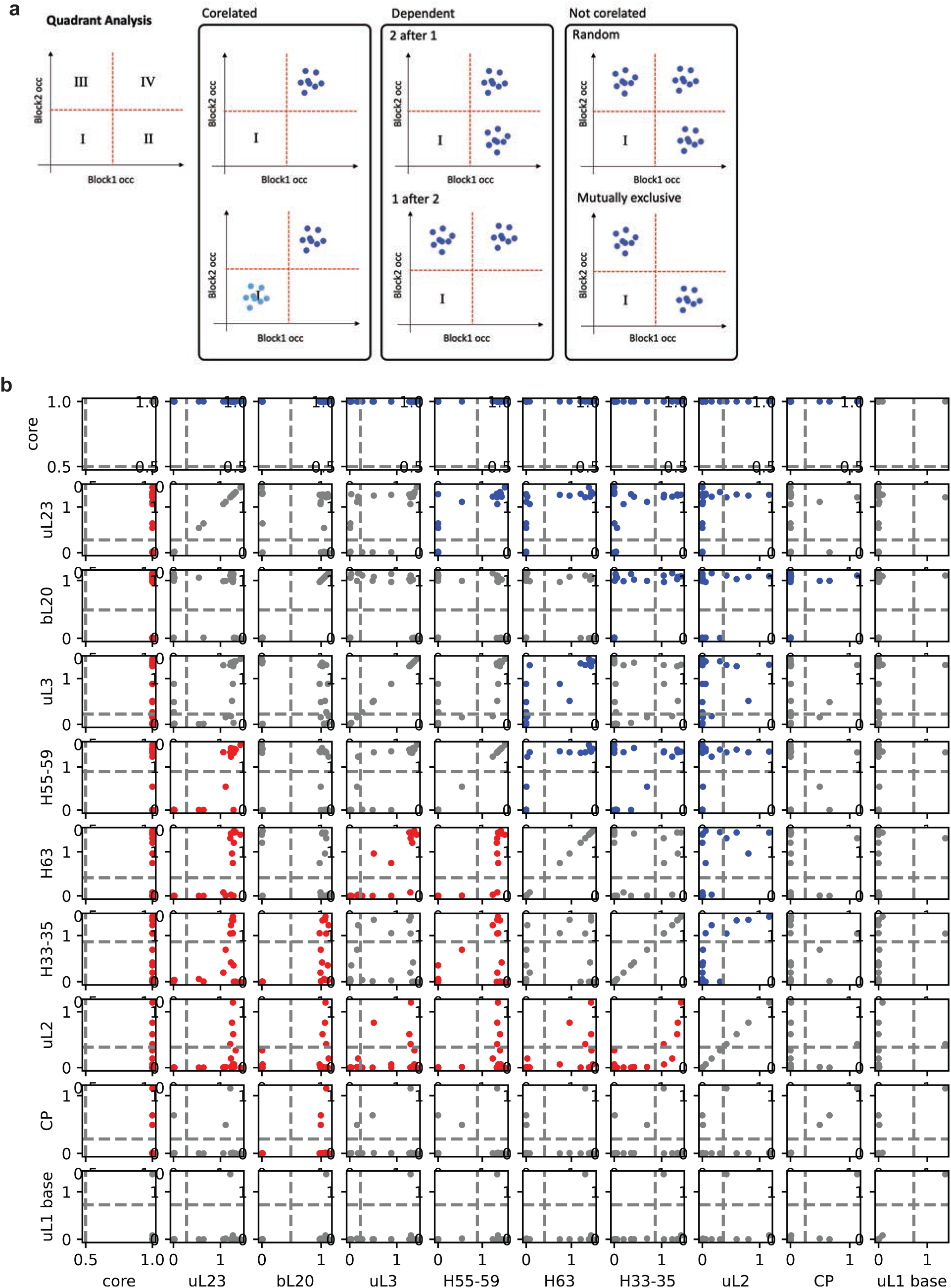
Quadrant analysis of assembly blocks of ΔdeaD dataset. **a**) Description of block dependency relationships with a quadrant plot. The scatter plot of pairwise block occupancy is divided into four quadrants (Q-I, II, III and IV). Each point represents a density map. For example, points in Q-II represent an intermediate map where block 1 is folded (occupied) while block 2 not folded (not occupied). The plots fall into three major categories, which are correlated, dependent, and not correlated. **b**) The pairwise quadrant analysis of 10 assembly blocks from ΔdeaD dataset is displayed. Plots showing a dependency of block 1 on block 2 are colored in red, and plots showing a depdency of block 2 on block 1 are shown in blue, while uncorrelated plots are shown in gray. (See Methods for details)

**Extended data Fig. 5.**
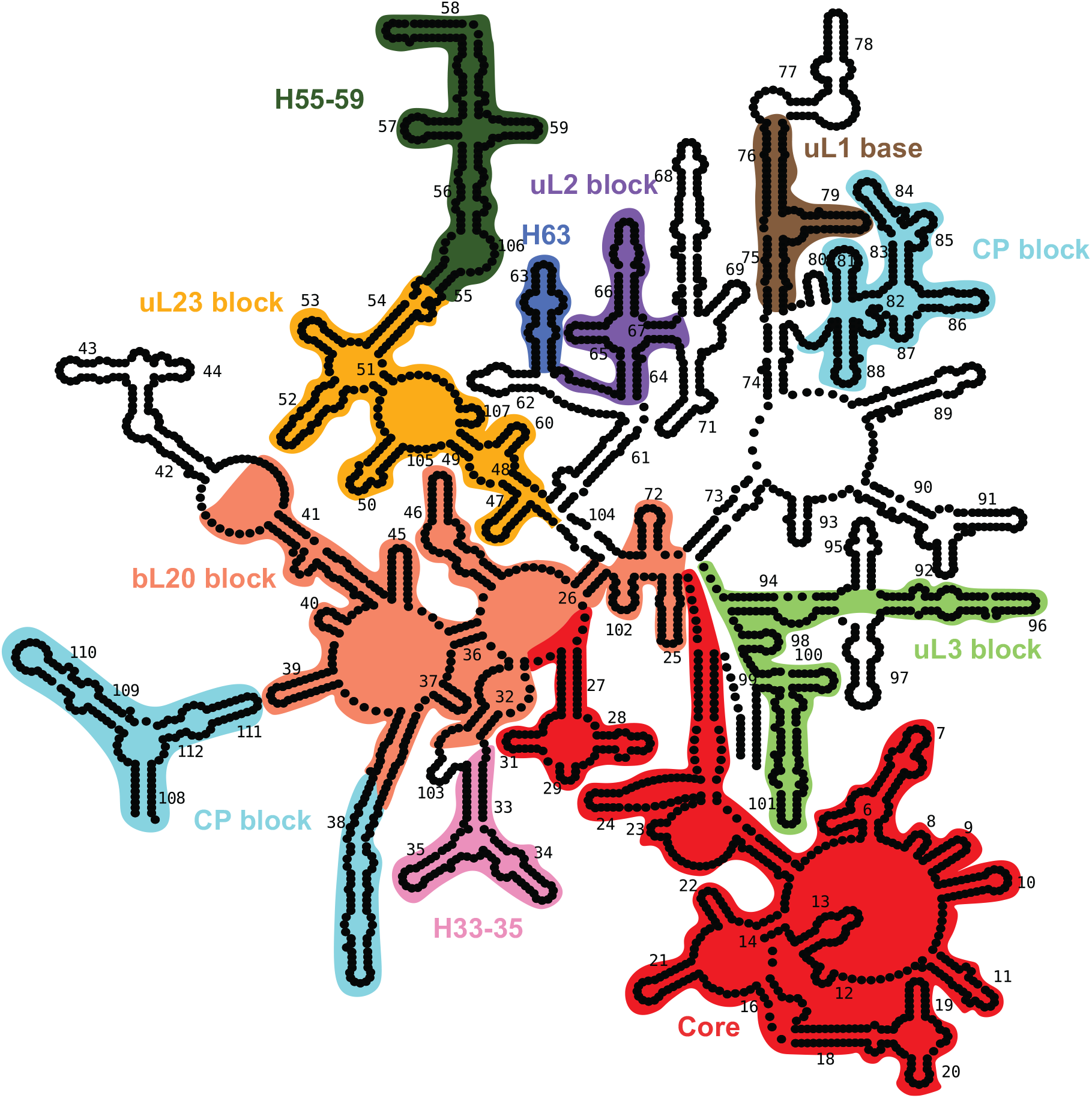
Assembly blocks displayed on the 23S/5S Secondary structure. The rRNA helices are colored according to the assembly blocks shown in Fig. 2 and Fig 3b.

**Extended data Fig. 6.**
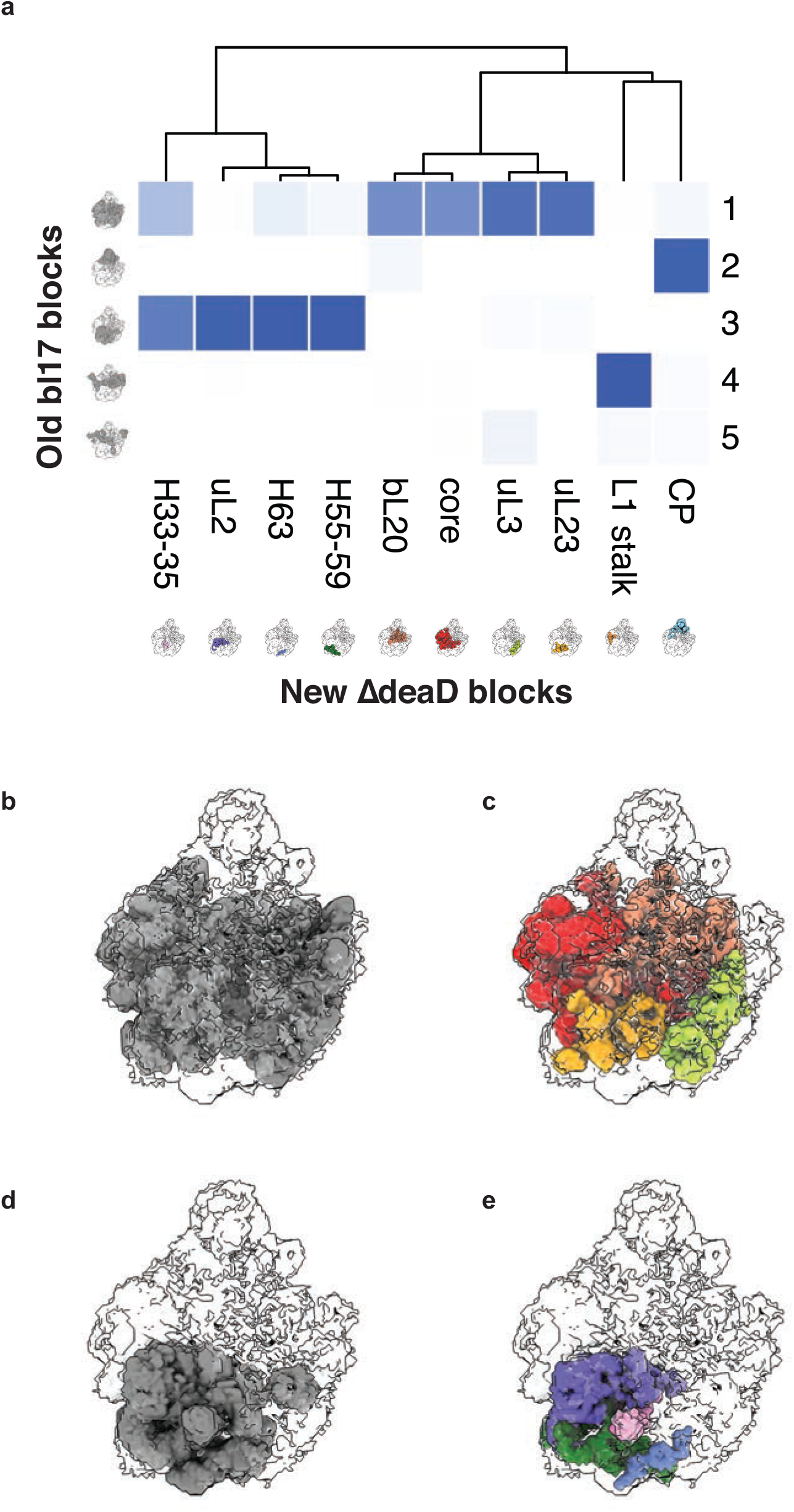
Subdivision of Previous bL17 blocks. **a**) Voxel occupancy map of new blocks in previous bL17 blocks. ##REF. Display of the 3D volumes shows that the previous bL17 old block 1 (**b**) can be subdivided into the assembly core, bL20, uL23 and uL3 blocks (**c**). Similarly, the previous bL17 block 3 (**d**) can be subdivided into the H55-58, H33-35, H63 and uL2 blocks (**e**).

**Extended data Fig. 7.**
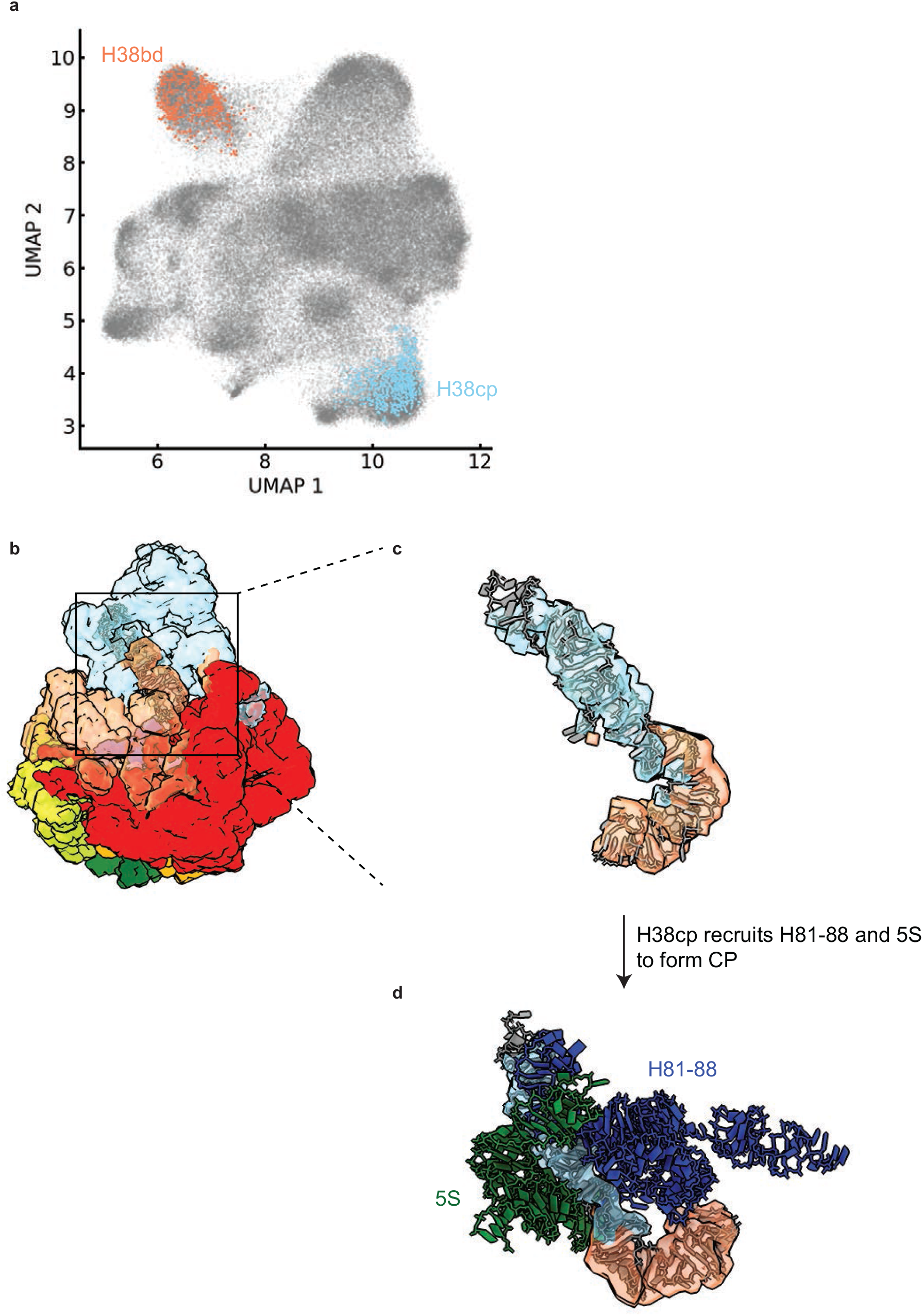
Subdivision of H38 led to insights into CP formation. **a**) Scatter plot of UMAP1/2 with H38bd and H38cp colored according to their respective blocks, showing clean separation of voxels. **b**) The atomic model of H38 from 4YBB (grey) was aligned to the assembly blocks (colored according to Fig. 2), resulted in subdivision of H38 into a body part (H38bd) as part of the bL20 block and CP part (H38cp) as part of the CP block (**c**). (**d**) The atomic model of 5S (green) and H81-88 (dark blue) are displayed, showing close contact with H38cp.

**Extended data Fig. 8.**
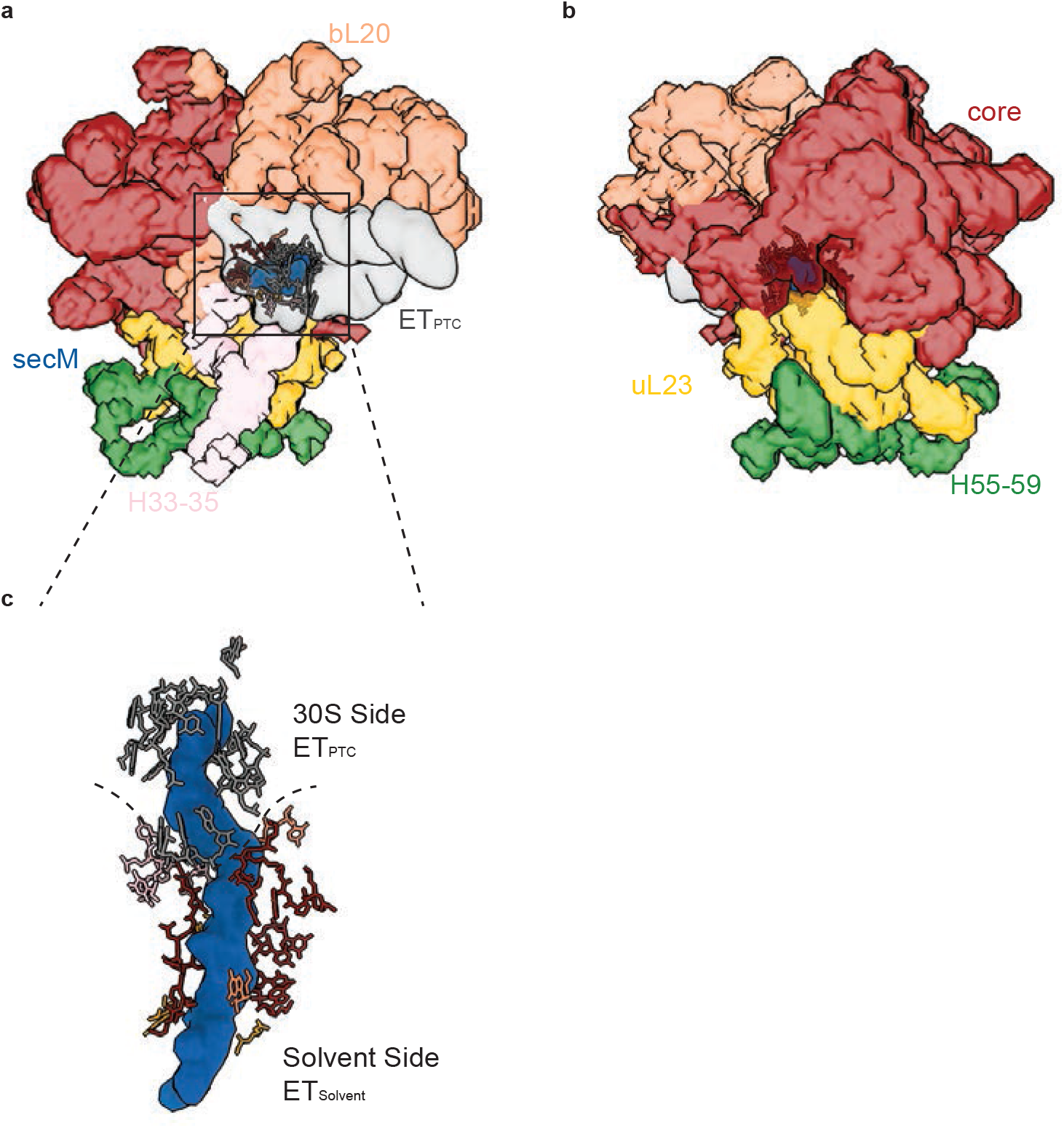
ET_solvent_ and ET_PTC_. **a-b**) Blocks surrounding the peptide exit tunnel are shown, with direct contact residues in stick form. The ETsolvent blocks are colored according to Fig. 2, the ET_PTC_ is colored in grey, and the SecM nascent chain density (molmap from PDB: 3JBU) is colored in dark blue. **c**) Contacting residues are displayed around the SecM nascent chain using same color coding in **a-b**).

**Extended data Fig. 9.**
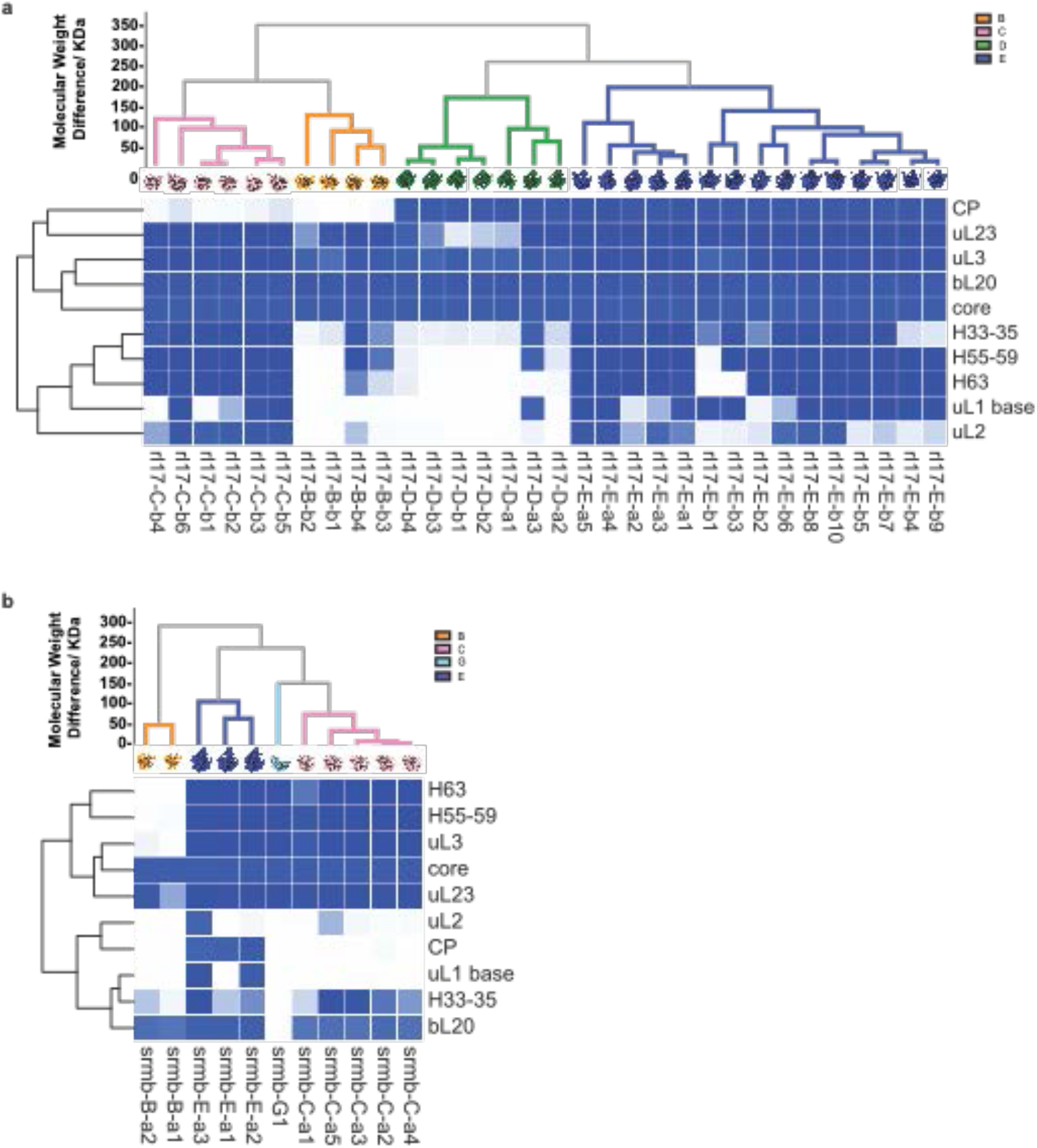
Comparison of intermediates from ΔsrmB and bL17 depletion with blocks from the DdeaD dataset. The density maps from previous studies were analyzed as an occupancy matrix based on the assembly blocks from the DdeaD dataset, clustering the matrix along both axes. The dendrograms are colored according to the major classes, B,C,D, E, and G. (**a**) bL17-depletion (**b**) ΔsrmB datasets.

**Extended data Fig. 10.**
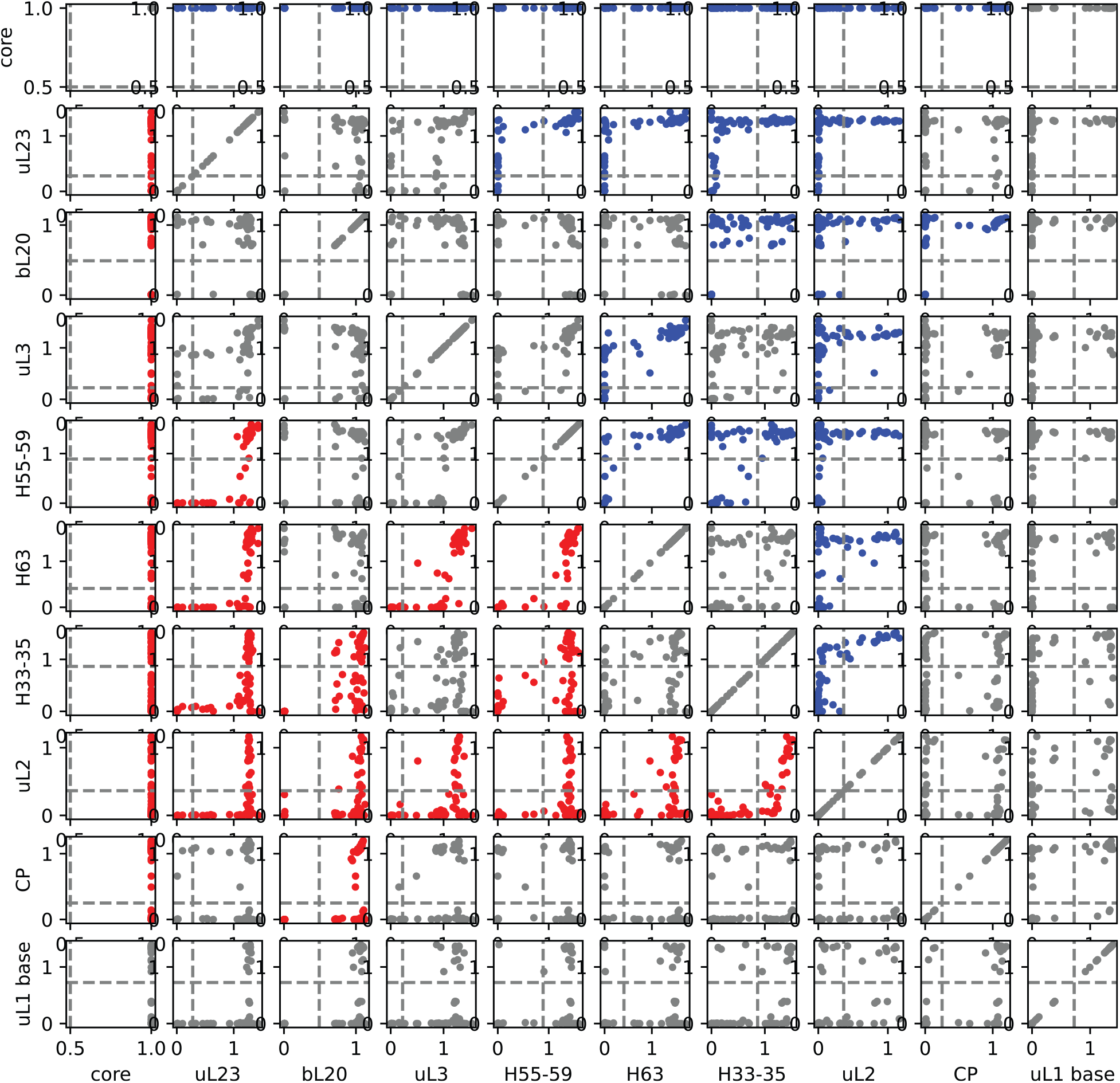
Quadrant analysis to determine the dependency of assembly blocks for three datasets. Pairwise quadrant analysis of the 10 assembly blocks from the ΔdeaD dataset on three datasets (64 maps) is displayed. Color coding is same as Extended data Fig. 4.

**Extended data Fig. 11.**
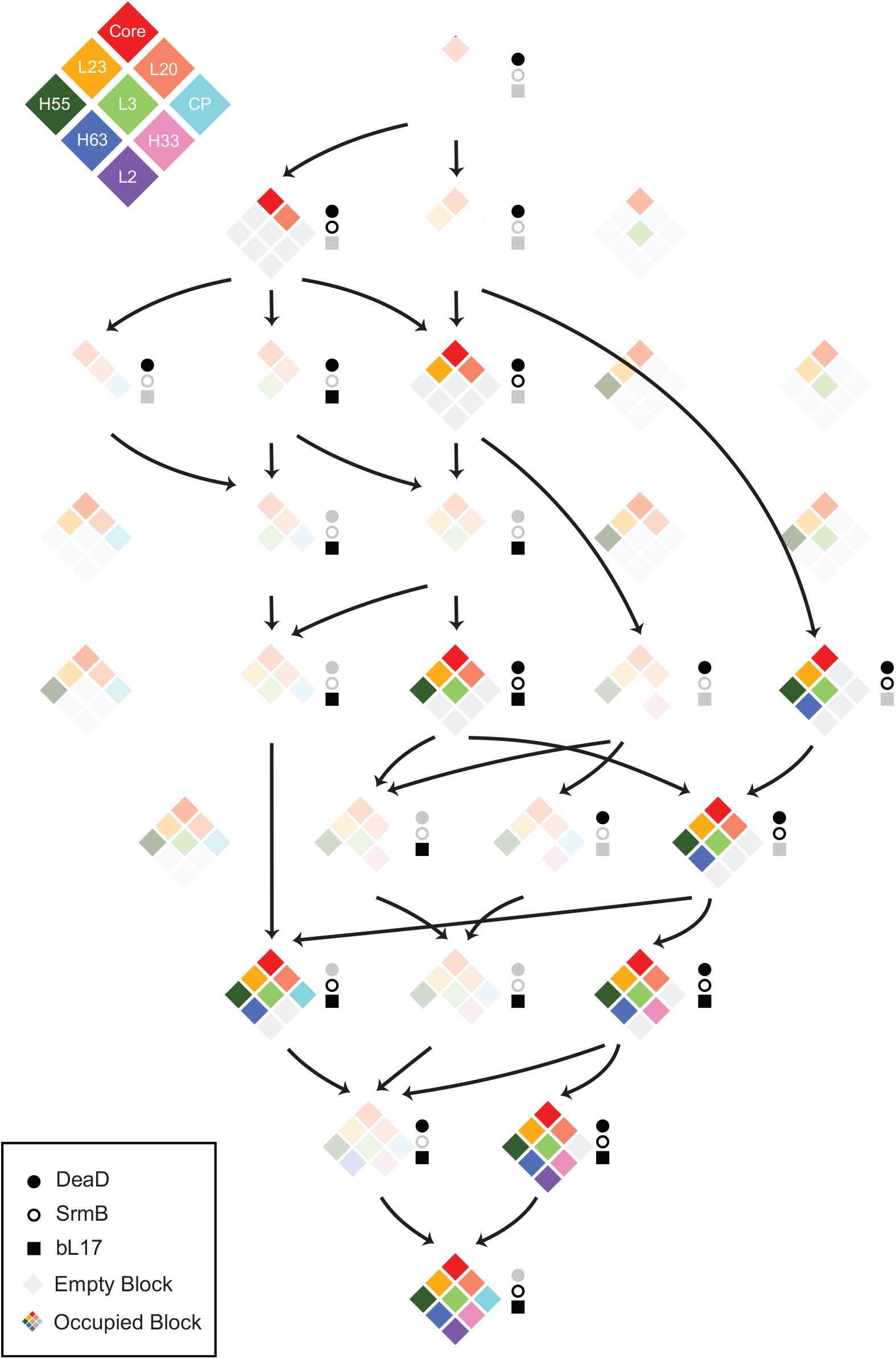
Diamond block assembly diagram for the ΔsrmB intermediates. All intermediate found in ΔsrmB are highlighted, with color coding the same as in Fig. 4.

**Extended data Fig. 12.**
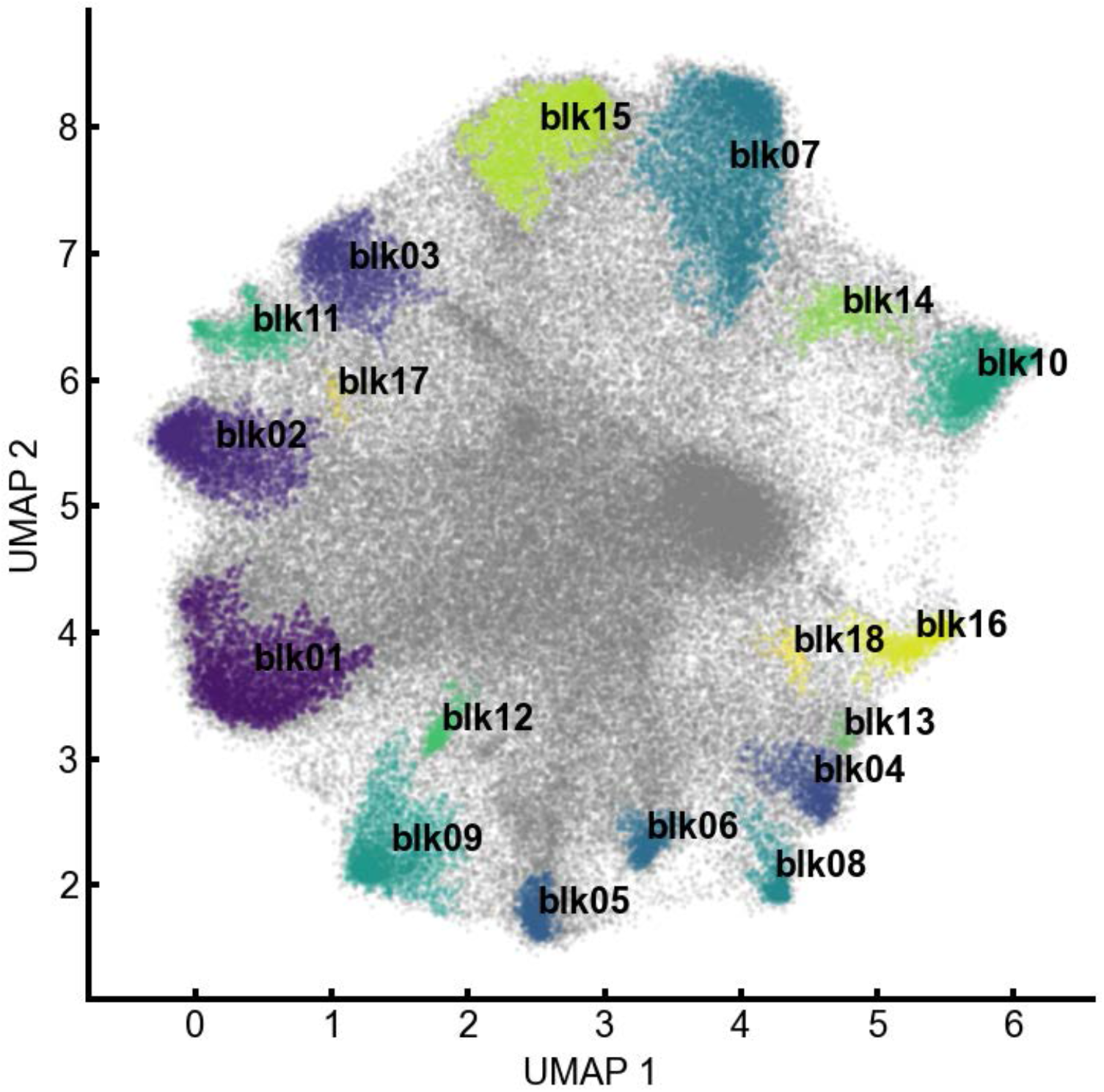
18 blocks PCA-UMAP-HDBSCAN. The PCA-UMAP-HDBSCAN segmentation was calculated using all three datasets, ΔdeaD, bL17, and, ΔsrmB. Voxels above a 1% threshold in three datasets are well organized in the newly calculated UMAP space, with 18 contiguous volume blocks extracted by clustering with HDBSCAN colored. Extraction of features from the PCA-UMAP-HDBSCAN representation is straightforward.

**Extended data Fig. 13.**
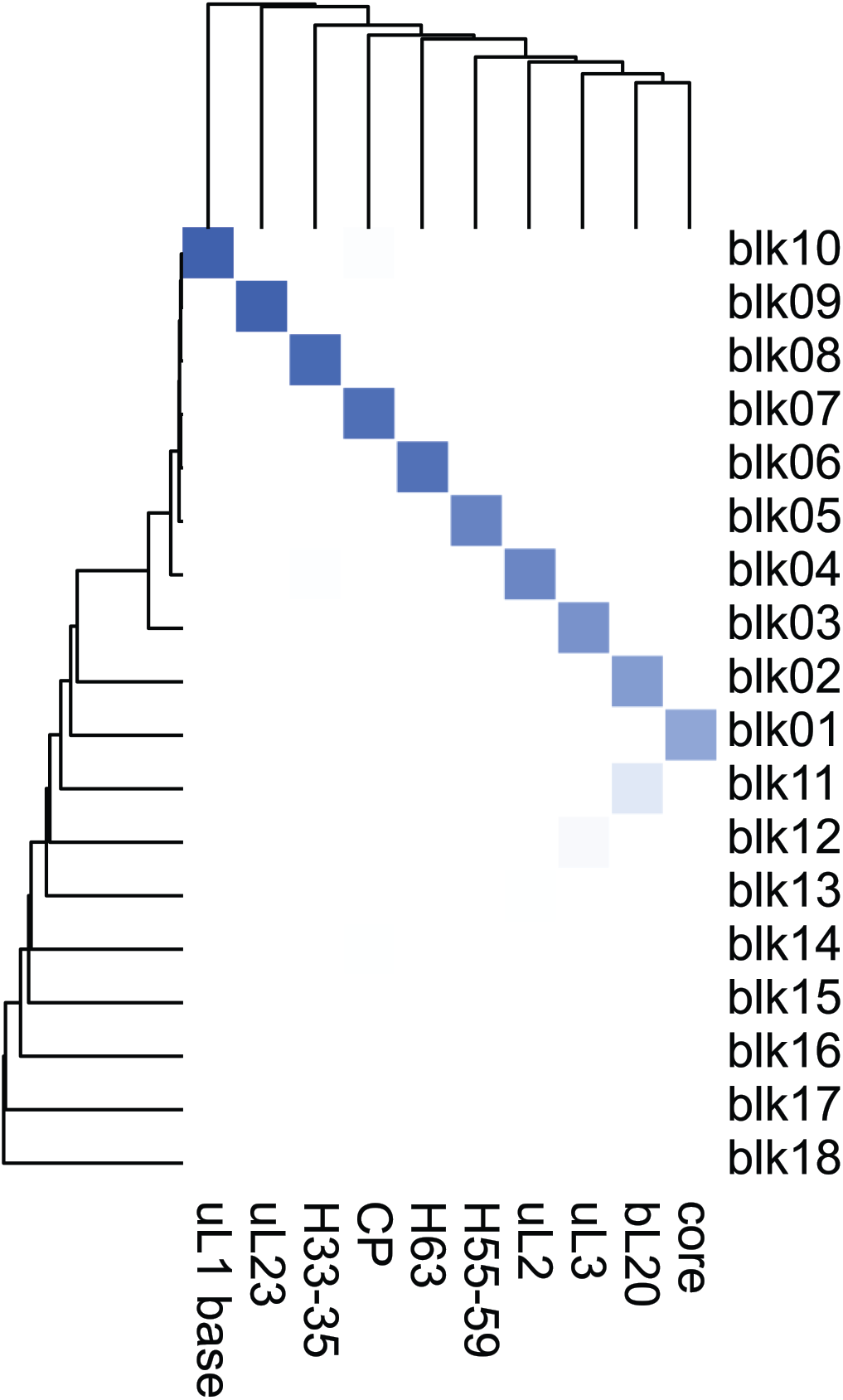
Occupancy analysis of 18 assembly blocks derived from all three datasets. Voxel occupancy matrix for the new segmentation with 18 blocks in terms of the 10 blocks from ΔdeaD dataset, and displayed with clustering of both dimensions using Euclidean distance with Ward linkage. The extracted features are very similar, with some subdivision of the smaller set of blocks.

**Extended data Fig. 14.**
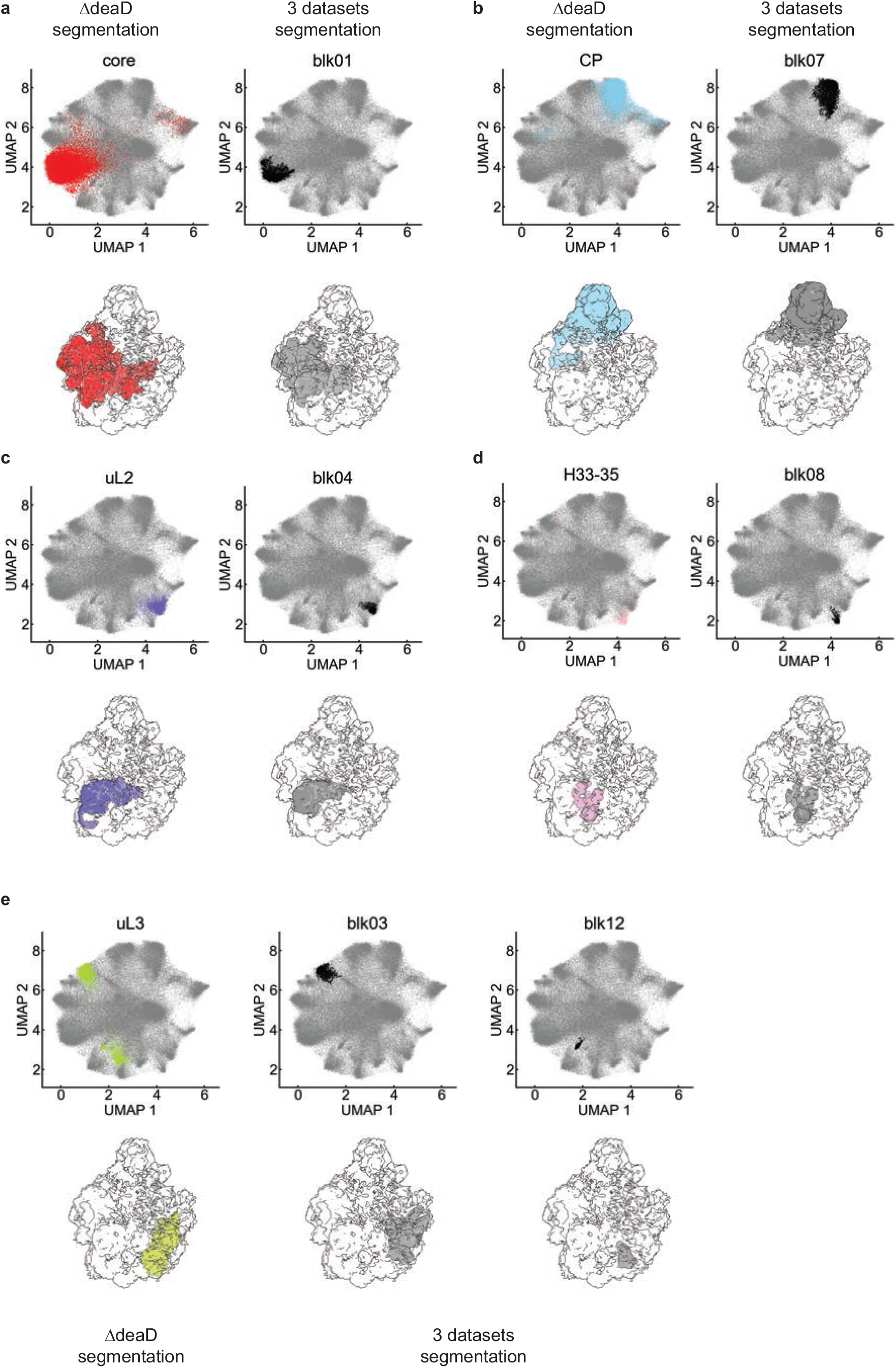
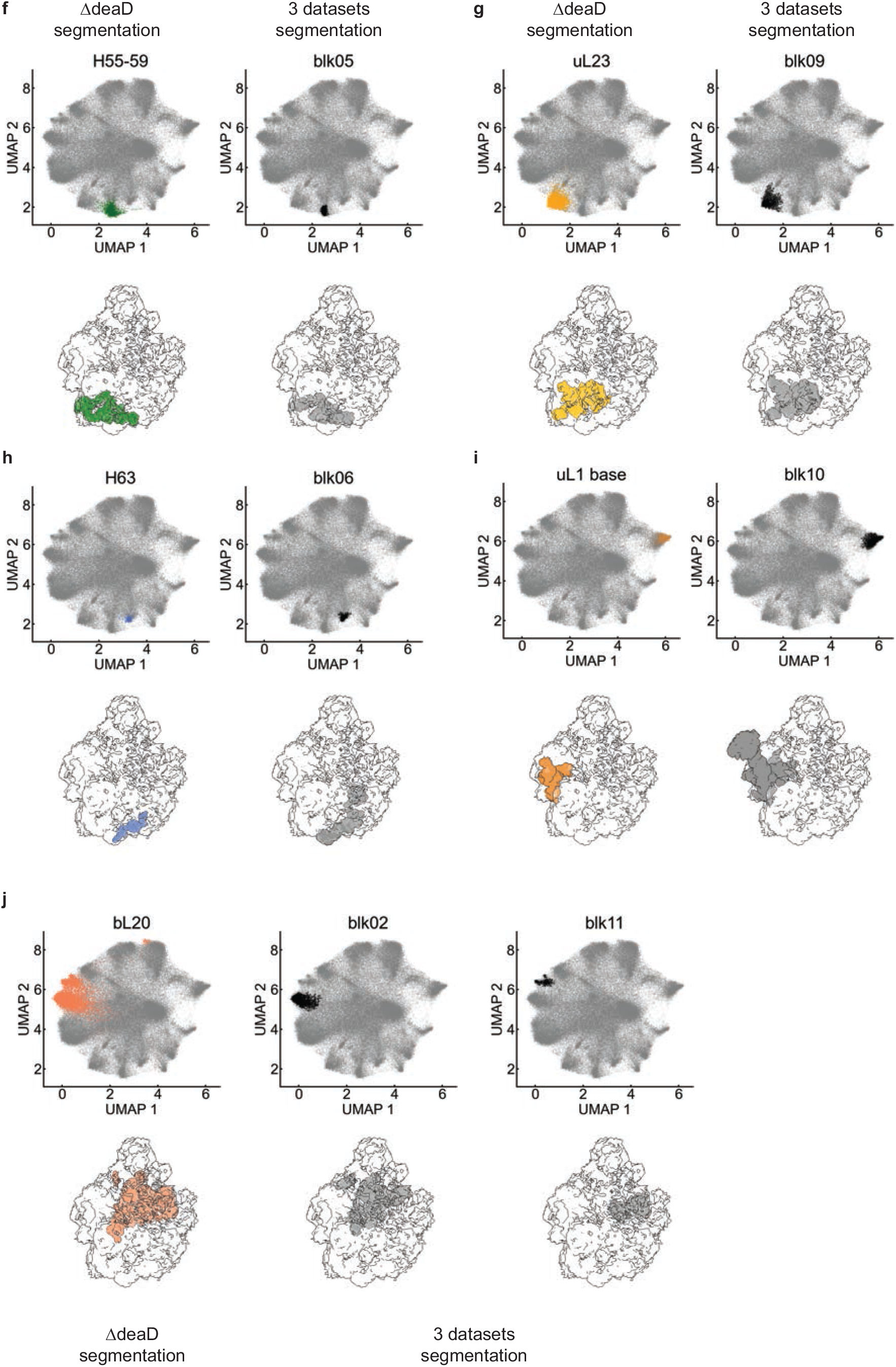
Comparison of 18 blocks from PCA-UMAP-HDBSCAN analysis of three datasets with 10 blocks from the DdeaD dataset. Among the 18 blocks obtained from three datasets, 12 blocks are corresponding to 10 blocks from the ΔdeaD dataset alone. For each comparison (part I/II a-d), on the top left, ΔdeaD blocks are colored according to Fig. 2, while corresponding three dataset blocks are on the top right colored in black in UMAP1/2 scatterplots. the corresponding 3D volumes outlined in the 1% threshold 50S, with ΔdeaD blocks colored according to Fig.2 and the new 18 blocks colored in grey. Notably, for Part I/II e, two cases of subdivision of a ΔdeaD block are displayed with the same color coding as above.

**Extended data Fig. 15.**
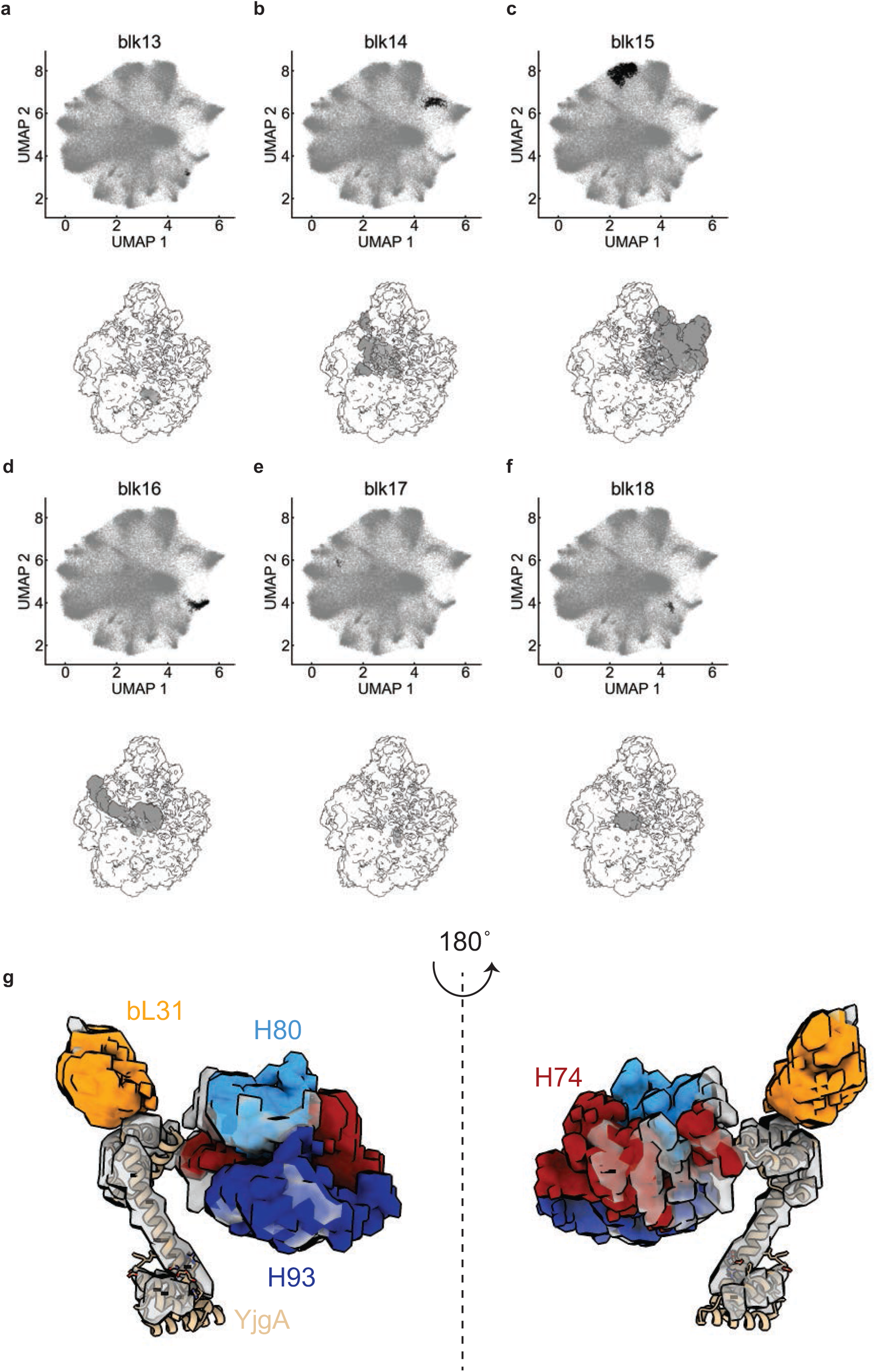
Six new blocks identified from the PCA-UMAP-HDBSCAN analysis of three datasets. **a-f**) display six new blocks in three dataset PCA-UMAP-HDBSCAN, both in UMAP space and as 3D volumes. The composition of the volumes in terms of 50S helices and proteins are described in the SI. g) Display of the volume corresponding to non-native density, identified as yjgA in blk14 (transparent grey), aligned to the atomic model of yjgA in yellow. Density for bL31, H74, H80 and H93 volumes from 4YBB are also shown in solid colored volume. ***is this all one new block?

**Extended data Fig. 16.**
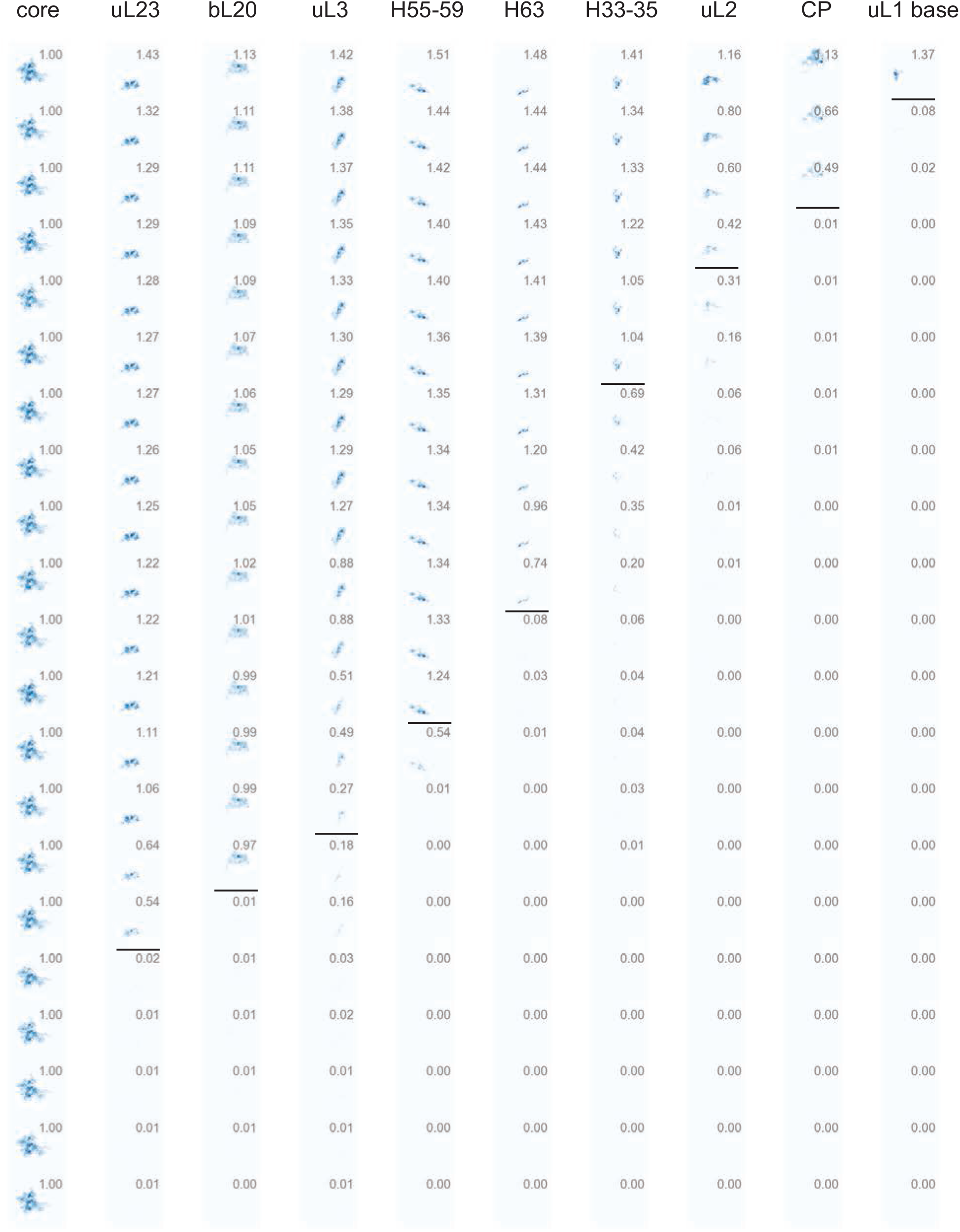
Thresholding the presence of 10 blocks in the set of 21 ΔdeaD density maps. Each column shows a heatmap of the set of 21 ΔdeaD maps after masking by one certain block, ordered from largest occupancy to smallest normalized occupancy of the sblock displayed near the heatmap. The thresholding line is drawn in black for each block.

## Notes

### Competing Interest Statement

The authors have declared no competing interest.

