## Supplementary Material for "Unsupervised Voxel-based Segmentation reveals a Landscape of Bacterial Ribosome Large Subunit Early Assembly"

**Supplementary Information**

Table of contents

**1 Characterization of the strain and intermediate peaks**
1-1 pre50S intermediate accumulated in ΔdeaD strain upon cold shock
1-2 Mass spectrometry methods
1-2-1 Doubling time of WT and ΔdeaD strain
1-2-2 Whole cell proteomic analysis of ΔdeaD strain upon cold shock
1-2-3 SWATH mass spectrometry analysis of pre-50S/50S fractions
1-2-4 Quantitative rRNA modification analysis of pre-50S/50S fractions
**2 Workflow of Iterative classification with ab initio reconstruction in CryoSPARC**
2-1 Example in ΔdeaD workflow
2-2 Job history in CryoSPARC
2-3 Metadata table for refined maps
**3 Parameters in PCA-UMAP and method comparison**
3-1 Direct UMAP and different metrics
3-2 PCA-UMAP with different number of principle components
3-3 Nearest neighbor in UMAP
3-4 Summary table for PCA-UMAP
**4 Detailed ΔdeaD block description**
4-1 Assembly core
4-2 bL20 blocks
4-3 other blocks
**5 Dependency analysis**
5-1 Thresholding and Outlier curation
**6 rRNA modification**
**7 Detailed descriptions of three dataset block**
**8 Supplementary Information reference**

**1-1 pre50S intermediate accumulated in ΔdeaD strain upon cold shock**

DeaD is a cold shock protein in E. coli, with several annotated functions involving mRNA stability and association with the 50S ribosomal subunit [1-3]. Indeed, whole cell proteomics shows DeaD is induced 4-fold upon cold shock in wild type BW25113 strain, while the abundance of most of the ribosomal proteins remains unchanged (Figure S1). The ΔdeaD strain shows a severe growth defect at 19 C, but not at 37 C (Table S1), and the sucrose gradient of total cell lysate shows a decrease in the abundance of 70S ribosomes, an accumulation of free 30S subunits and new intermediate peaks arising between 30S and 50S (Figure S2), supporting the accumulation of pre-50S subunits, consistent with previous results [1]. Both the growth defect and sucrose gradient profile of the deletion strain can be rescued after introducing a DeaD expression plasmid, which further supports that the assembly defect comes from absence of DeaD. The effect could be either a direct effect on 50S assembly, or indirect due to an effect of the DEAD-box in regulating some ribosomal protein expression level ***REF. Whole cell proteomics on the deletion strain at 19 C, clearly shows that there is no significant alteration of r-protein levels, which implies more direct impact of DeaD on ribosome biogenesis. The intermediate fraction was also analyzed by quantitative protein mass spectrometry, showing partial depletion of bL35, bL36, bL28, bL32, bL34, bL33, uL30 and uL6 proteins (Figure S3).

**1-2 Mass spectrometry methods**

**1-2-1 Doubling time of WT, ΔdeaD and ΔdeaD-pHSL-deaD strain**

WT, ΔdeaD and ΔdeaD-pHSL-deaD strains were inoculated in LB medium and grown overnight, then diluted into fresh 500 mL LB (1:200) at 19 or 37 $^{o}$C. Either 0 or 2.5 nM HSL was added into the ΔdeaD-pHSL-deaD strain during cell culture. The $OD_{600}$ was monitored during growth until mid-log stage. The time-coursed $OD_{600}$ was used to fit the doubling time in the expression for exponential growth in GraphPad Prism using non-linear regression. Cells from the WT and ΔdeaD culture harvested at $OD_{600}$ = 0.3-0.5 by centrifugation, and prepared for sucrose gradient analysis (see Methods) and mass spectrometry analysis.

**1-2-2 Whole cell proteomic analysis**

WT cells were grown in $^{15}$N M9 medium at 37 ˚C as $^{15}$N spike for quantitation. For ΔdeaD cells grown at 19 and 37 ˚C in as described in section 1-2-1, 0.5 $OD_{600}$ units of cell culture was was mixed with 0.5 $OD_{600}$ units of $^{15}$N spike culture, and the cells were pelleted by centrifugation at 14,000 rpm for 30 seconds. The cell pellet was washed with cold water for twice, before 13% TCA was added directly to the pellet, followed by vortex mixing and incubation for 12 hours at 4$^{o}$C. The protein precipitate was pelleted by centrifugation at 14,000 rpm for 30 min at 4$^{o}$C. The protein pellet was washed twice by adding 700 ul 10% TCA followed by centrifugation at 14,000 rpm. The final supernatant was removed, and the pellet was dried in a Speed-Vac concentrator, and then dissolved in 40 ul of fresh 100 mM ammonium bicarbonate and 5% acetonitrile (ACN). To break the disulfide bonds between cysteine residues, a 4 ul aliquot of 50 mM DTT was added to each sample, and the samples were incubated at 65 ˚C for 10 min. Thiol groups were blocked by the addition of 4 ul of 100 mM iodoacetamide (IAA) and incubation for 30 min at 30 ˚C in the dark. The protein was digested by the addition of 2 ul of 0.1 ug/ul modified sequence grade porcine trypsin (Promega, Co., Madison, WI) with incubation overnight at 37 ˚C. Undigested proteins were removed by adding 1/3 volume of 2% trifluoroacetic acid in 20% ACN with

centrifugation, and the supernatant was transferred to a Pierce C-18 spin column (Thermo Fisher Scientific Inc., Rockford, IL) to remove salts and to concentrated the peptides. The sample was eluted from the column with 60 ul 70% ACN in 0.1% formic acid. The eluant was dried in a Speed-Vac concentrator and the peptides were resuspended in 10 ul 5% ACN in 0.1% formic acid.
Peptides were analyzed on a 5600+ Triple TOF mass spectrometer (T-TOF) coupled to Eksigent ekspert™ nanoLC 400 systems with ChromXP C18 (3µm 120 A) column. The sample was injected via autosampler into the column with a trap-eluting mode using NanoLC Trap ChromXP C18 (3µm 120 A). NanoLC separation consisted of the following steps: (1) a linear gradient from 10% to 40% buffer B over 80 min, (2) column washing with 80% B for 20 min, and (3) column equilibration with 10% B for 30 min. The MS/MS data was collected with data-dependent acquisition mode on the T-TOF. The data was pre-processed according to Trans-Proteomic Pipeline[4]. Briefly, the raw data was transformed with MSConverter. Peptides were identified using X!Tandem[5] searching against custom E. coli database (derived from UniProt organism 83333) and combined into spectral libraries using SpectraST[6] before relative quantification in Massacre as previously described [7, 8]. The Massacre fitting results were filtered based on an in-house random forest filtering algorithm trained with peptide Massacre fits that were manually annotated as good or bad.

#### 1-2-3 SWATH mass spectrometry analysis of pre-50S/50S fractions

For SWATH sample preparation, 50 pmol of $^{15}$N WT-70S was added to pre-50S fractions that were purified by sucrose gradient centrifugation. TCA was added to the spiked fraction to 13% final concentration to precipitate proteins. The tryptic peptide preparation was described in section 1-2-2. After resuspension, peptides were analyzed on the T-TOF coupled to an Agilent 1100 Series HPLC instrument with capillary flow electrospray (Agilent Technologies Inc., Santa Clara, CA). Peptides were injected using an autosampler onto an Agilent Zorbax SB C18 150 mm x 0.5 mm HPLC column, with the mobile phase 0.1% formic acid in HPLC grade H2O and acetonitrile with 0.1% formic acid. SWATH data-independent mass spectrometry data were collected using positive polarity over the m/z range of 300-1300. The mass spectral data were analyzed using Skyline [9], and the r-protein levels were quantified as previously described [10].

#### 1-2-4 Quantitative rRNA modification analysis of pre-50S/50S fractions

The rRNA in pre-50S and 70S fractions were extracted with Direct-zol™ RNA MiniPrep Kits (Zymo) using the manufacturer's instructions. The RNA MS/MS was performed as previously described [11]. The rRNA isolated from mature $^{15}$N-ribosomes was added as an external standard. Purified $^{14}$N 23S from the test sample and $^{15}$N-RNA spike were mixed in approximately 1:1 molar ratio, heat-denatured, and digested with RNase T1 or A for 1 h at 55 °C in 25 mM ammonium acetate (pH = 6). Nucleolytic fragments were separated on an Xbridge C18 column (Waters) using buffer A (15 mM ammonium acetate, pH = 8.8) and buffer B (15 mM ammonium acetate, pH = 8.8 and 50% acetonitrile). HPLC separation consisted of the following steps: (1) isocratic elution with 1% buffer B for 5 min, (2) a linear gradient from 1% to 15% buffer B over 40 min, (3) column washing with 100% B for 25 min, and (4) column equilibration with 1% B for 30 min. Data were recorded over the 400–1700 m/z range on Agilent Q-TOF 6520 ESI. Pairs of the co-eluting $^{14}$N- and $^{15}$N-labeled fragments were identified using Pytheas [12]. After the LC-MS peak profiles were extracted and fit using Agilent MassHunter.

### 2 Workflow of Iterative classification with ab initio reconstruction in CryoSPARC

A screenshot of the cryoSPARC work flow for reconstruction of intermediate density maps in the ΔdeaD dataset is shown in Figure S4. The overall workflow is hierarchical, with the outputs of one job serving as the inputs to a subsequent series of jobs. As described in Methods, the cleaned particle stacks after RELION preprocessing were submitted to direct *ab initio* reconstruction, asking for 4 classes in job J62, shown in Figure S5a. Classes 0 and 3 did not have interpretable particles, while classes 1 and 2 showed clear shapes of putative 50S intermediates (Figure S5a-d). The 4 classes from J62 were analyzed by 2D classification (J168-171, Figure S5e-h) to validate our picks for next round. These two stacks were subjected to the next round of *ab inito* reconstruction, again with 4 classes, in jobs J63 and J64. In general, the subclassification was continued iteratively, as shown in Figure S4, until one of two termination criteria was satisfied. As one example, the particle numbers in the subclassification job J130 were 1763, 32, 84 and 1742 for subclasses, 0,1,2, and 3, respectively, (green box in Figure S4).. Classes 0 and 3 were below the particle threshold of 2000, and termination was affected by 3D refinement in jobs J147 and J148, followed by *ab initio* reconstruction and 3D refinement. As a second example, after jobs J136 and J137, a 3D refinement of two subclasses yielded maps with differences below the 10 KDa threshold, so the particles were merged and subjected to *ab initio* reconstruction with 1 class (J283, red box in Figure S4). Finally, for terminal species, *ab initio* reconstruction for 1 class (J283) was performed with 3D refinement (NEW) job (J304), before performing ThreeDFSC (job J326) for FSC calculation and resolution estimation (Purple box in Figure S4). Class metadata for three datasets can be found in Table S2 and the detailed information of iterative classification can be found in the Table S4.

### 3 Parameters in PCA-UMAP and method comparison

This section discusses the optimization process of segmentation of a set of electron density maps using the dimensionality reduction methods of PCA and UMAP. The goal is to identify groups of voxels with similar behavior across the set of maps using a clustering method, and tuning the parameters of the PCA-UMAP to optimize performance and generation of interpretable segments. For scatter plot in this section, blocks are colored according to Figure 2 (the final optimal PCA-UMAP-HDBSCAN segmentation) in the main text, to illustrate the quality of block separation for subsequent cluster picking.

#### 3-1 Direct UMAP and different metrics

The PCA-UMAP results described in Methods was compared to the direct application of UMAP on the 21 x 114,392 matrix without PCA transformation. Also, four metrics (Euclidean, Canberra, correlation and cosine) were applied in both the direct UMAP method or PCA-UMAP method, for comparison (Figure S6). We found that two UMAP-1,2 scatter plots are the same for direct UMAP and PCA-UMAP using the Euclidean metric. This may be expected since the PCA is a matrix factorization, which should not change the Euclidean distance if all PCs are used. (Figure S6 b f). However, the blocks are better-defined in the PCA-UMAP compared to direct UMAP using the other three metrics (Canberra, correlation, and cosine). Among the eight different trials, PCA-UMAP with Canberra metric showed separated blocks with smallest average distance to cluster median with smallest variation (Figure S7).

Since, UMAP is inherently stochastic[13], the stability of the algorithm was tested by running the same dataset for 100 times using direct UMAP and PCA-UMAP with Canberra metric with different seeds. The mean squared displacement (MSD) for each dot in the set of scatter plots is calculated. The MSD scatter plot (Figure S8a) showed that the algorithm for PCA-UMAP with Canberra metric was very stable and a given block does not vary significantly between runs. In contrast direct application of UMAP exhibited highly variable positions of the blocks. (Figure S8b)

**3-2 PCA-UMAP with different number of principle components**

When UMAP was applied to PCA transformed features, we can control the number of features input to UMAP. Using a low number of PCs, the UMAP step was unable to separate clusters. With increasing number of PCs, the UMAP step stabilizes, such that there is little additional change when additional PCs are included. The Euclidean metric led to convergence with a smaller number of PCs, but the resulting distribution of blocks was elongated and tangled, making the subsequent extraction process difficult (Figure S9). Using the Canberra metric, gave a more separable distribution. (Figure S10). Both the Canberra and Euclidean metrics showed convergence with increasing PC number, suggesting that the higher PCs mostly represent the noise (Extended Data Fig.2), having little influence on cluster formation. In practice, we use the full PCA reconstruction to reduce the possibility of subjective feature picking.

**3-4 Nearest neighbors in UMAP**

Figure S11 showed the tuning of the parameter for number of nearest neighbors in UMAP. We found if the number becomes too large, some clusters began to merge in to the large blocks. On the other hand, if the number of nearest neighbors is too small, the model captured some details that do not represent a whole block. We chose an intermediate value with 100 nearest neighbors, which roughly corresponds to the 3 KDa volume, corresponding approximately to a small RNA helix.

**3-5 Summary table for PCA-UMAP**

| Method | Direct PCA | Direct UMAP | PCA-UMAP(Euclidean) | PCA-UMAP(Canberra) |
|---|---|---|---|---|
| Cluster identification | No | Yes | Yes | Yes |
| Cluster extraction | No | No | spectra distribution | Yes |
| Reproducibility | Same every time | block jumping (Canberra) | Not tested | Good |

**4 Detailed ΔdeaD block description**

**4-1 Assembly core**

The first block common to all maps in the dataset is termed the **Assembly core** as the origin of the assembly pathway. It consists of all helices in 23S Domain I, except H1, and four helices (H27-29, H31) in Domain II 23S rRNA, and the four early r-proteins, uL4, uL22, uL24 and uL29. uL24 binds to the most 5′ end is reported to be one of the two initiators for 50S formation in *in vitro* experiment [14]. Although uL22 has contacts with all domains of 23S rRNA, the structured region of domain I rRNA (~600 nucleotides) is sufficient for uL22 to stably bind to 23S rRNA. This implies that uL22 can associate early with the nascent transcript. Interestingly, the Assembly core does not include H1 which is formed by the base pairing of 5′ and 3′ of the entire 23S rRNA, proximal to the RNase cleavage site that liberates the pre-50S particle from the primary rRNA transcript. This is the first evidence showing the early 50S assembly intermediates do not need H1 formation, which is strong evidence for the co-transcriptional block-wise assembly.

**4-2 bL20 Block**

The next largest block is the **bL20 Block,** which contains uL13, bL20, bL21, bL34 and most of the helices in domain II of 23S rRNA, with exception for four helices in assembly core, H33-35 and H42-44. The latter helices H42-44 correspond to the base for the right stalk where uL10/11 anchor which is not present in any of the intermediates in this data set. This block also contains a portion of the long helix 38 (H38bd) (Extended Data Fig.7), while the remaining portion has more interaction with the central protuberance (H38cp).

**4-3 other blocks**

The **uL23 Block** contains uL23 and majority of domain III of 23S rRNA, except h55-59 and h106. The **uL3 block** contains uL3, bL17, bL19 and domain VI 23S of rRNA. Protein uL3 was the second initiator protein identified based on *in vitro* reconstitution experiments [14]. The CP block contains 5S rRNA, the distal portion of H38 (H38cp), H81-88, and proteins L5, L15, L18, L25, L27, and L30. The uL2 block contains uL2 and part of domain IV rRNA. Other than these RNA-protein blocks, there are also three small blocks solely consisting of RNA helices, which are the H55-59, H33-35, H63, and uL1 base block.

**5 Dependency analysis**

**5-1 Redundant edge pruning**

Figure S12 showed criteria of removing an edge in the dependency network. Briefly, if block 2 depends on block 1, at the same time, block 3 depends on both block 1 and block 2. To simplify the model, we can remove edge from block 1 to block 3, without changing the fact of block 3 needs formation of both block 1 and block 2.

**6 rRNA modification**

**H33-35 formation and implication of early-stage rRNA modification**

Within the H33-35 block, H35 is modified at positions $m^1G$ (745), Ψ (746), and $m^5U$ (747). Quantitative RNA mass spectrometry shows that double methylation in the pre-50S fraction is over 95%, (Figure S13, Table S5) while only about 40% of particles in the set of maps have H35 properly folded and docked. This evidence strongly suggests that the substrate for the corresponding enzymes RlmA and RlmC is a very immature pre50S particle. RlmA was shown to bind on free 23S[15], however, substrate for RlmC is still unknow. Our results implied the early function of these methyl transferase in ribosome biogenesis.

**7 Detailed descriptions of three dataset blocks**

| 18 blocks | contents |
|---|---|
| blk01 | Core block |
| blk02 | bl20 block – H25 uL13 |
| blk03 | uL3 block |
| blk04 | uL2 block |
| blk05 | H55-59 block |
| blk06 | H63 block |
| blk07 | CP block |
| blk08 | H34 |
| blk09 | uL23 |
| blk10 | uL1 base |
| blk11 | uL13 H25 |
| blk12 | bL17 |
| blk13 | H62 |
| blk14 | H74 93 80 bL33 yjgA |
| blk15 | uL10/11 stalk |
| blk16 | H68-70 |
| blk17 | H104 |
| blk18 | uL14 |

### 8 Supplementary Information reference

10. Davis, J.H., et al., *Modular Assembly of the Bacterial Large Ribosomal Subunit.* Cell, 2016. **167**(6): p. 1610-1622 e15.
11. Popova, A.M. and J.R. Williamson, *Quantitative analysis of rRNA modifications using stable isotope labeling and mass spectrometry.* J Am Chem Soc, 2014. **136**(5): p. 2058-69.
12. D'Ascenzo, L., et al., *Pytheas: a software package for the automated analysis of RNA sequences and modifications via tandem mass spectrometry.* Nat Commun, 2022. **13**(1): p. 2424.
13. McInnes, L., J. Healy, and J. Melville, *Umap: Uniform manifold approximation and projection for dimension reduction.* arXiv preprint arXiv:1802.03426, 2018.
14. Nowotny, V. and K.H. Nierhaus, *Initiator proteins for the assembly of the 50S subunit from Escherichia coli ribosomes.* Proc Natl Acad Sci U S A, 1982. **79**(23): p. 7238-42.
15. Hansen, L.H., F. Kirpekar, and S. Douthwaite, *Recognition of nucleotide G745 in 23 S ribosomal RNA by the rrmA methyltransferase.* J Mol Biol, 2001. **310**(5): p. 1001-10.

**Table S1** Doubling time of different strain at 19 ˚C

| Strain | Plasmid | Medium | HSL conc. (nM) | Doubling Time (min)* |
|---|---|---|---|---|
| BW25113 | - | LB | 0 | 122 |
| ΔdeaD | - | LB | 0 | 370 |
| ΔdeaD | pHSL-deaD* | LB | 0 | 163 |
| ΔdeaD | pHSL-deaD | LB | 0.1 | 177 |
| ΔdeaD | pHSL-deaD | LB | 0.5 | 199 |
| ΔdeaD | pHSL-deaD | LB | 1 | 189 |

* pHSL-*deaD* plamid is constructed from pHSL-*rplQ*, replacing *rplQ* with *deaD*.
* * Doubling time was fitted in Prism using non-linear regression (exponential growth) Single colony was picked from LB agar plates and inoculate in LB medium at 37 $^{o}$C overnight. The next day, overnight culture was diluted (1:200) into 500 mL LB. The growing curve was recorded at 19 ˚C with vigorously shaking.

**Table S2** intermediate density map metadata

| Job ID | Class | Subclass | Corrected Resolution | Particle Number | Voxel | In dataset name | Name | Project ID |
|---|---|---|---|---|---|---|---|---|
| 307 | B | B-a | 5.91 | 2490 | 22043 | B-a1 | dead-B-a1 | 11 |
| 309 | B | B-a | 5.41 | 3303 | 22847 | B-a2 | dead-B-a2 | 11 |
| 308 | B | B-a | 5.62 | 3125 | 22867 | B-a3 | dead-B-a3 | 11 |
| 311 | B | B-a | 5.41 | 8604 | 25437 | B-a4 | dead-B-a4 | 11 |
| 312 | B | B-a | 6.24 | 1911 | 28430 | B-a5 | dead-B-a5 | 11 |
| 310 | B | B-a | 5.99 | 1387 | 31601 | B-a6 | dead-B-a6 | 11 |
| 313 | B | B-b | 6.43 | 1689 | 27444 | B-b1 | dead-B-b1 | 11 |
| 299 | C | C-a | 6.14 | 2028 | 31141 | C-a1 | dead-C-a1 | 11 |
| 300 | C | C-a | 5.96 | 2112 | 32584 | C-a2 | dead-C-a2 | 11 |
| 301 | C | C-a | 6.16 | 1763 | 33412 | C-a3 | dead-C-a3 | 11 |
| 302 | C | C-a | 5.41 | 6810 | 33905 | C-a4 | dead-C-a4 | 11 |
| 298 | C | C-a | 6.27 | 1500 | 36134 | C-a5 | dead-C-a5 | 11 |
| 303 | C | C-b | 5.41 | 2416 | 41387 | C-b1 | dead-C-b1 | 11 |
| 304 | C | C-b | 5.41 | 3338 | 37531 | C-b2 | dead-C-b2 | 11 |
| 293 | E | E-a | 5.43 | 2312 | 50393 | E-a1 | dead-E-a1 | 11 |
| 297 | G | G | 6.74 | 2017 | 25794 | G1 | dead-G1 | 11 |
| 296 | G | G | 5.89 | 5464 | 26579 | G2 | dead-G2 | 11 |
| 294 | G | G | 5.41 | 5758 | 29195 | G3 | dead-G3 | 11 |
| 295 | G | G | 5.74 | 3019 | 30902 | G4 | dead-G4 | 11 |
| 306 | B | preB | 6.39 | 2837 | 13990 | preB1 | dead-preB1 | 11 |
| 314 | B | preB | 7.36 | 1103 | 16696 | preB2 | dead-preB2 | 11 |
| 1831 | B | B-b | 5.45 | 2685 | 30703 | B-b1 | rl17-B-b1 | 26 |
| 1830 | B | B-b | 6.00 | 1606 | 31052 | B-b2 | rl17-B-b2 | 26 |
| 1832 | B | B-b | 5.45 | 1772 | 32554 | B-b3 | rl17-B-b3 | 26 |
| 1833 | B | B-b | 5.45 | 1585 | 35814 | B-b4 | rl17-B-b4 | 26 |
| 1814 | C | C-b | 5.45 | 3068 | 39005 | C-b1 | rl17-C-b1 | 26 |
| 1815 | C | C-b | 5.45 | 2943 | 39801 | C-b2 | rl17-C-b2 | 26 |
| 1811 | C | C-b | 5.45 | 4368 | 41135 | C-b3 | rl17-C-b3 | 26 |
| 1813 | C | C-b | 5.54 | 1277 | 41342 | C-b4 | rl17-C-b4 | 26 |
| 1812 | C | C-b | 5.45 | 3438 | 43118 | C-b5 | rl17-C-b5 | 26 |
| 1810 | C | C-b | 5.45 | 4655 | 45238 | C-b6 | rl17-C-b6 | 26 |
| 1801 | D | D-a | 5.45 | 2084 | 37456 | D-a1 | rl17-D-a1 | 26 |
| 1802 | D | D-a | 5.45 | 2377 | 40580 | D-a2 | rl17-D-a2 | 26 |
| 1803 | D | D-a | 5.45 | 1964 | 42170 | D-a3 | rl17-D-a3 | 26 |
| 1804 | D | D-b | 5.45 | 1651 | 42152 | D-b1 | rl17-D-b1 | 26 |
| 1805 | D | D-b | 5.45 | 1914 | 42857 | D-b2 | rl17-D-b2 | 26 |
| 1806 | D | D-b | 5.45 | 1582 | 43879 | D-b3 | rl17-D-b3 | 26 |
| 1807 | D | D-b | 5.45 | 1506 | 45093 | D-b4 | rl17-D-b4 | 26 |
| 1819 | E | E-a | 5.45 | 2468 | 47104 | E-a1 | rl17-E-a1 | 26 |
| 1821 | E | E-a | 5.45 | 1554 | 47445 | E-a2 | rl17-E-a2 | 26 |
| 1820 | E | E-a | 5.45 | 1921 | 48373 | E-a3 | rl17-E-a3 | 26 |
| 1835 | E | E-a | 5.45 | 11791 | 48927 | E-a4 | rl17-E-a4 | 26 |
| 1816 | E | E-a | 5.45 | 13258 | 52135 | E-a5 | rl17-E-a5 | 26 |
| 1809 | E | E-b | 5.45 | 6390 | 47629 | E-b1 | rl17-E-b1 | 26 |
| 1829 | E | E-b | 5.45 | 1949 | 55635 | E-b10 | rl17-E-b10 | 26 |
| 1822 | E | E-b | 5.68 | 1273 | 49021 | E-b2 | rl17-E-b2 | 26 |
| 1808 | E | E-b | 5.45 | 1532 | 50798 | E-b3 | rl17-E-b3 | 26 |
| 1825 | E | E-b | 5.45 | 1968 | 52095 | E-b4 | rl17-E-b4 | 26 |
| 1823 | E | E-b | 5.45 | 1467 | 52913 | E-b5 | rl17-E-b5 | 26 |
| 1827 | E | E-b | 5.45 | 1428 | 53397 | E-b6 | rl17-E-b6 | 26 |
| 1824 | E | E-b | 5.45 | 1892 | 53474 | E-b7 | rl17-E-b7 | 26 |
| 1828 | E | E-b | 5.45 | 1532 | 54563 | E-b8 | rl17-E-b8 | 26 |
| 1826 | E | E-b | 5.45 | 1818 | 55005 | E-b9 | rl17-E-b9 | 26 |
| 202 | B | B-a | 9.27 | 1234 | 19913 | B-a1 | srmb-B-a1 | 15 |
| 190 | B | B-a | 8.29 | 1419 | 24089 | B-a2 | srmb-B-a2 | 15 |
| 196 | C | C-a | 7.75 | 4355 | 24078 | C-a1 | srmb-C-a1 | 15 |
| 197 | C | C-a | 8.08 | 1787 | 29870 | C-a2 | srmb-C-a2 | 15 |

| | | | | | | | | |
|---|---|---|---|---|---|---|---|---|
| 204 | C | C-a | 6.81 | 7185 | 30332 | C-a3 | srmb-C-a3 | 15 |
| 198 | C | C-a | 8.46 | 2439 | 31184 | C-a4 | srmb-C-a4 | 15 |
| 200 | C | C-a | 7.89 | 1525 | 35286 | C-a5 | srmb-C-a5 | 15 |
| 193 | E | E-a | 7.56 | 2485 | 36449 | E-a1 | srmb-E-a1 | 15 |
| 194 | E | E-a | 7.77 | 1624 | 41658 | E-a2 | srmb-E-a2 | 15 |
| 192 | E | E-a | 6.99 | 1827 | 43453 | E-a3 | srmb-E-a3 | 15 |
| 195 | G | G | 6.84 | 9918 | 24535 | G1 | srmb-G1 | 15 |

* Job or Project ID refer to CryoSPARC job or project ID.
* Classes were assigned according to Figure 1d.
* “Corrected resolution” was obtained from ThreeDFSC in Å.
* “Voxel” is voxel number above 1.00 threshold in the density map.

**Table S3** 5 Å residues around nascent peptide chain in 3JBU and 5NWY

| PDB | Entity | Site | contents | secondary structure | block |
|---|---|---|---|---|---|
| 3JBU | 23S | 460-461 | AC | H23 | core |
| | | 470 | A | H23 | |
| | | 507-508 | AA | H24 | |
| | | 752 | A | H35 | H33-35 |
| | | 790 | U | H32 | bL20 |
| | | 789 | A | H103 | |
| | | 1321 | A | H50 | uL23 |
| | | 1614 | A | H107 | |
| | | 2061-2063 | GAC | H74 | PTC |
| | | 2451-2452 | AC | H74/H89 | |
| | | 2505-2506 | GU | H90 | |
| | | 2585 | U | H90 | |
| | | 2609-2611 | UCC | H93/H73 | |
| | uL4 | 61 | R | | core |
| | uL22 | 82-85 | MKRI | | |
| | | 90-95 | KGRADA | | |
| | uL23 | 71-72 | GQ | | uL23 |
| 5NWY | 23S | 460-463 | ACCG | H23 | core |
| | | 470-471 | AA | H23 | |
| | | 746-752 | UUG--AA | H35 | H33-35 |
| | | 790-791 | UC | H32 | bL20 |
| | | 1259 | G | H26 | |
| | | 1323 | C | H50 | uL23 |
| | | 1614 | A | H107 | |
| | | 1782 | U | H65 | uL2 |
| | | 2057-2059 | GAA | H73/74 | PTC |
| | | 2061-2063 | GAC | H74 | |
| | | 2451 | A | H74/H89 | |
| | | 2505-2506 | GU | H90 | |
| | | 2581-2586 | G-GUUU | H90 | |
| | | 2602 | A | H93 | |
| | | 2609-2612 | UCCC | H93/H73 | |
| | uL4 | 57-67 | KKPWR-KGTGR | core | uL4 |
|  | uL22 | 83-95 | KRI----KGRADR | core | uL22 |

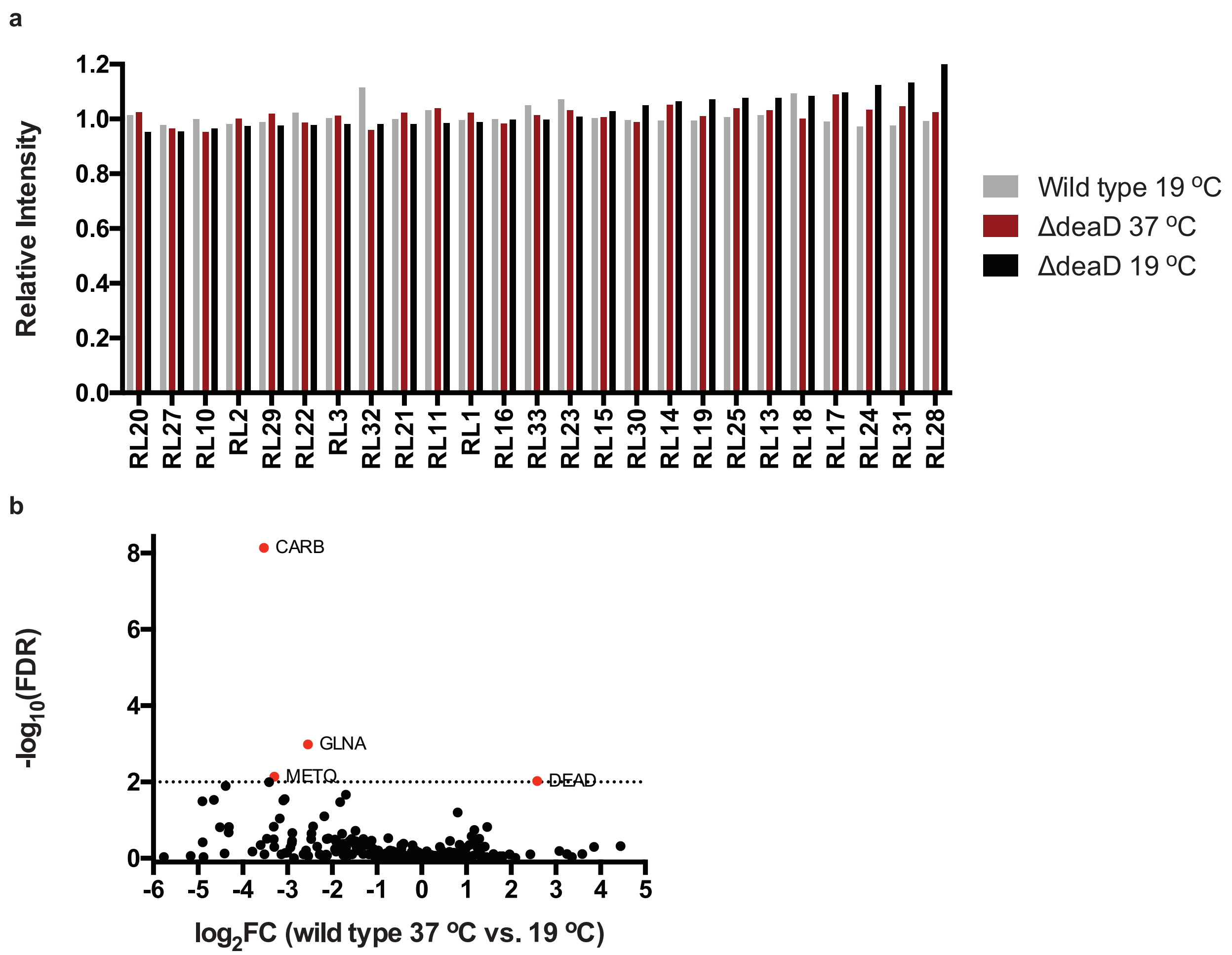


**Figure S1 | Proteomic analysis**

a) Quantitative mass spectrometry of large subunit ribosomal protein under various condition, intensity normalized to WT 37˚C. b) Volcano plot of wild type cells at 37 ˚C in M9 medium versus 19 ˚C in LB medium. Protein IDs above 1% FDR are colored in red.

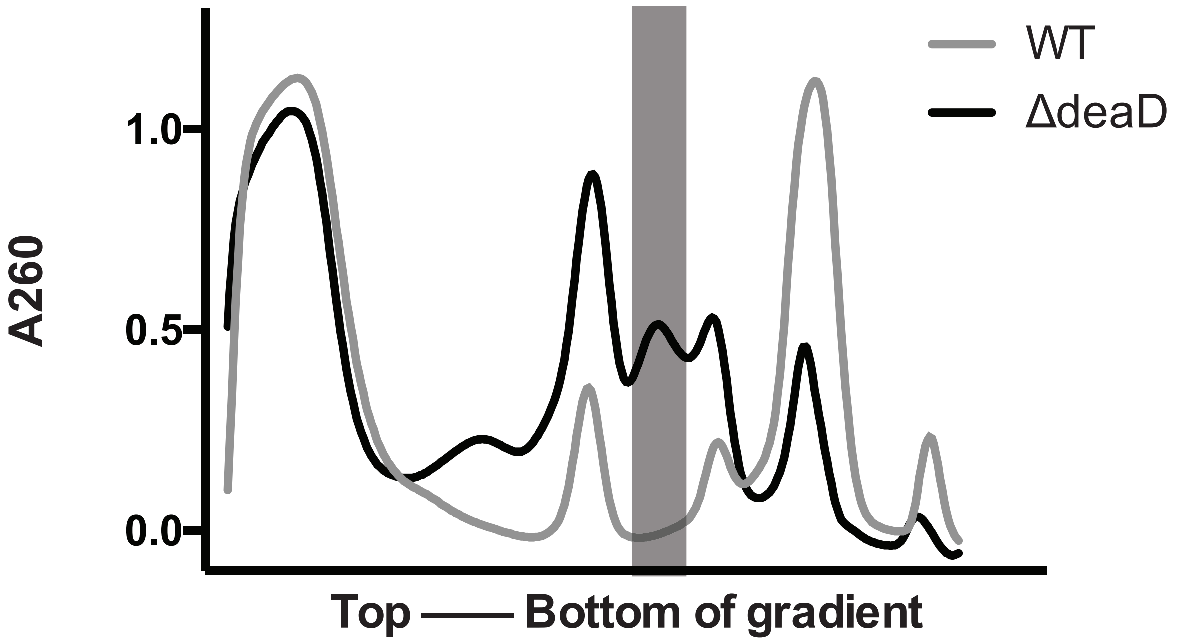


**Figure S2 | Sucrose gradient profiles showing distributions of ribosomal particles.**
10-40% Sucrose Gradient profile for cell lysates of WT (grey) and ΔdeaD (black) grown at 19 ˚C. The intermediate peak is shown in the grey transparent box.

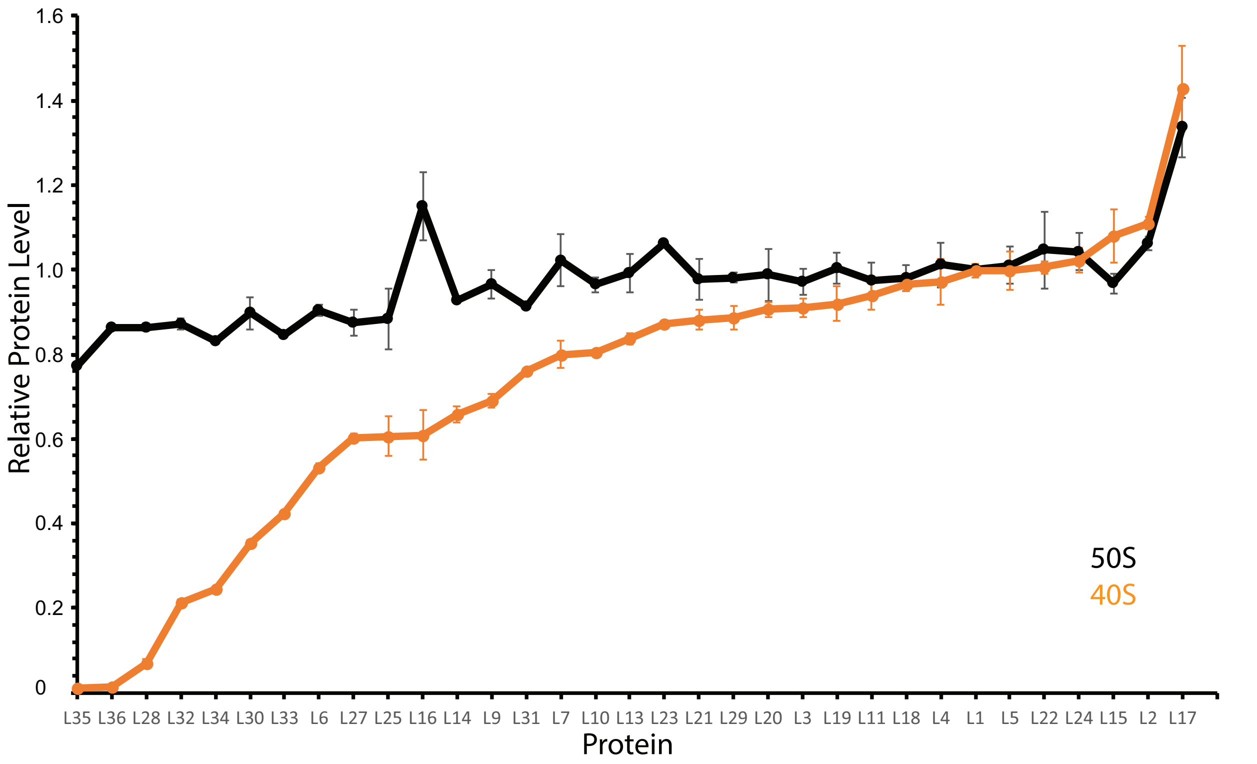


**Figure S3 | SWATH analysis of r-protein composition in gradient fractions**
SWATH quantification of large subunit ribosomal proteins in 40S fractions (orange) and 50S fractions (black) in the ΔdeaD strain.

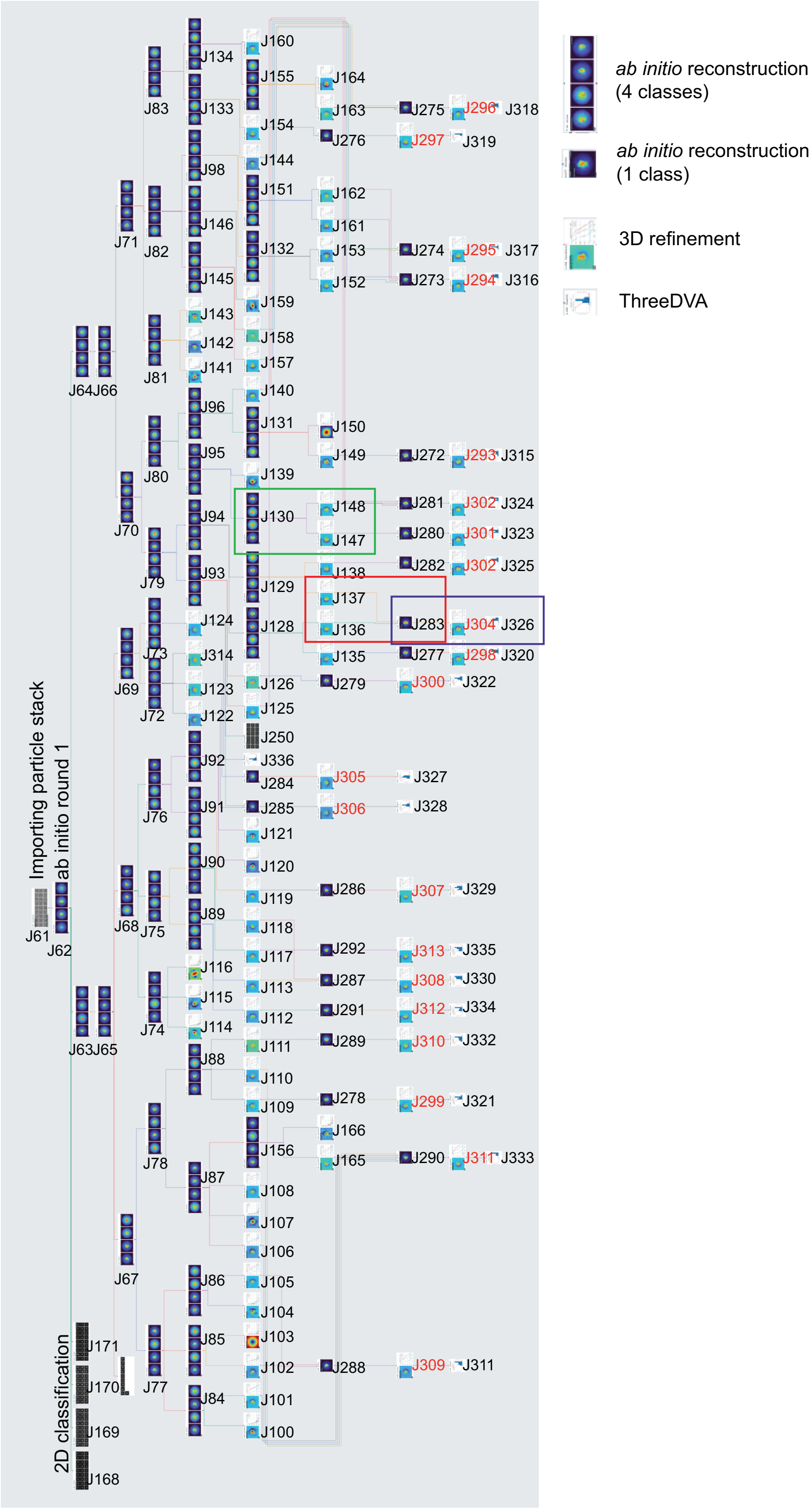

ab initio reconstruction (4 classes)
ab initio reconstruction (1 class)
3D refinement
ThreeDVA
Importing particle stack
ab initio round 1
2D classification
J61
J62
J63J65
J64J66
J67
J68
J69
J70
J71
J72
J73
J74
J75
J76
J77
J78
J79
J80
J81
J82
J83
J84
J85
J86
J87
J88
J89
J90
J91
J92
J93
J94
J95
J96
J98
J100
J101
J102
J103
J104
J105
J106
J107
J108
J109
J110
J111
J112
J113
J114
J115
J116
J117
J118
J119
J120
J121
J122
J123
J124
J125
J126
J128
J129
J130
J131
J132
J133
J134
J135
J136
J137
J138
J139
J140
J141
J142
J143
J144
J145
J146
J147
J148
J149
J150
J151
J152
J153
J154
J155
J156
J157
J158
J159
J160
J161
J162
J163
J164
J165
J166
J168
J169
J170
J171
J250
J272
J273
J274
J275
J276
J277
J278
J279
J280
J281
J282
J283
J284
J285
J286
J287
J288
J289
J290
J291
J292
J293
J294
J295
J296
J297
J298
J299
J300
J301
J302
J302
J304
J305
J306
J307
J308
J309
J310
J311
J312
J313
J314
J315
J316
J317
J318
J319
J320
J321
J322
J323
J324
J325
J326
J327
J328
J329
J330
J332
J333
J334
J335
J336
J311

**Figure S4 | Screenshot of CryoSPARC workflow for the ΔdeaD dataset analysis.**
A case of termination of subdivision using *ab initio* reconstruction highlighted with a green rectangle. An example of merging two similar classes that emerged in highlighted by a red rectangle. The final stage of 3D refinement and map validation by ThreeDFSC is highlighted by a purple rectangle.

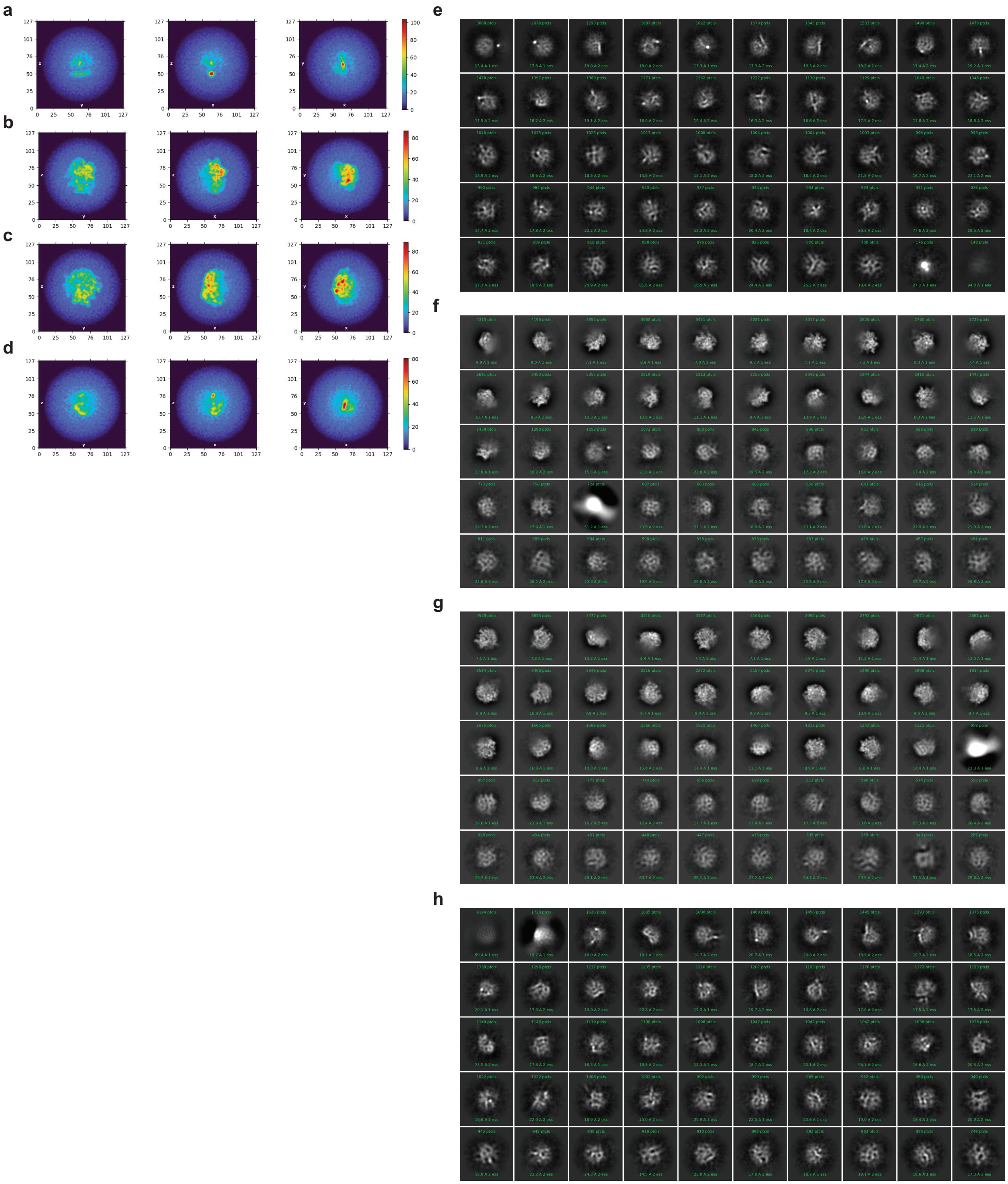

**Figure S5 | Examples of subclassification in analysis of ΔdeaD intermediates.**
**a-d)** Three perpendicular cross sections of the four initial classes of ab initio reconstruction (Job J62 in Figure S5). 2D classification for particles resulting from Job J62 from **e)** class 0, **f)** class 1 , **g)** class 2, and **h)** class 3.

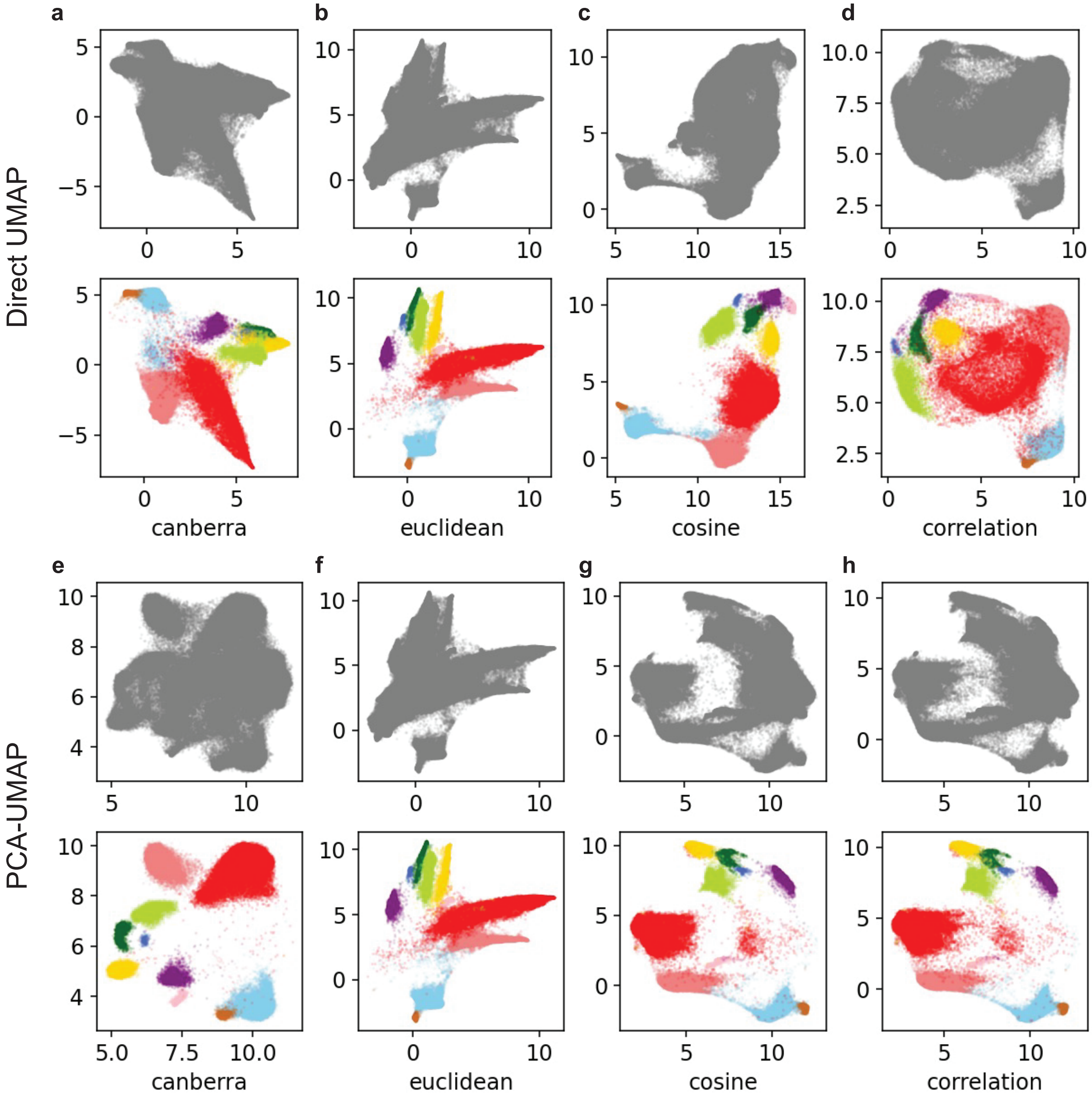


**Figure S6 | Comparison of direct UMAP and PCA-UMAP analysis using different metrics.** The ΔdeaD dataset was subjected to direct UMAP (**a-d**) or PCA-UMAP (**e-h**) of the set of voxels above threshold, with different metrics in UMAP (**a,e** for Canberra, **b,f** for Euclidean, **c,g** for cosine and **d,h** for correlation metric). For each approach, all voxels are colored in grey in the upper panel, and are colored according to the blocks assigned Fig.2 in the lower panel in corresponding UMAP1/2 space, for comparison.

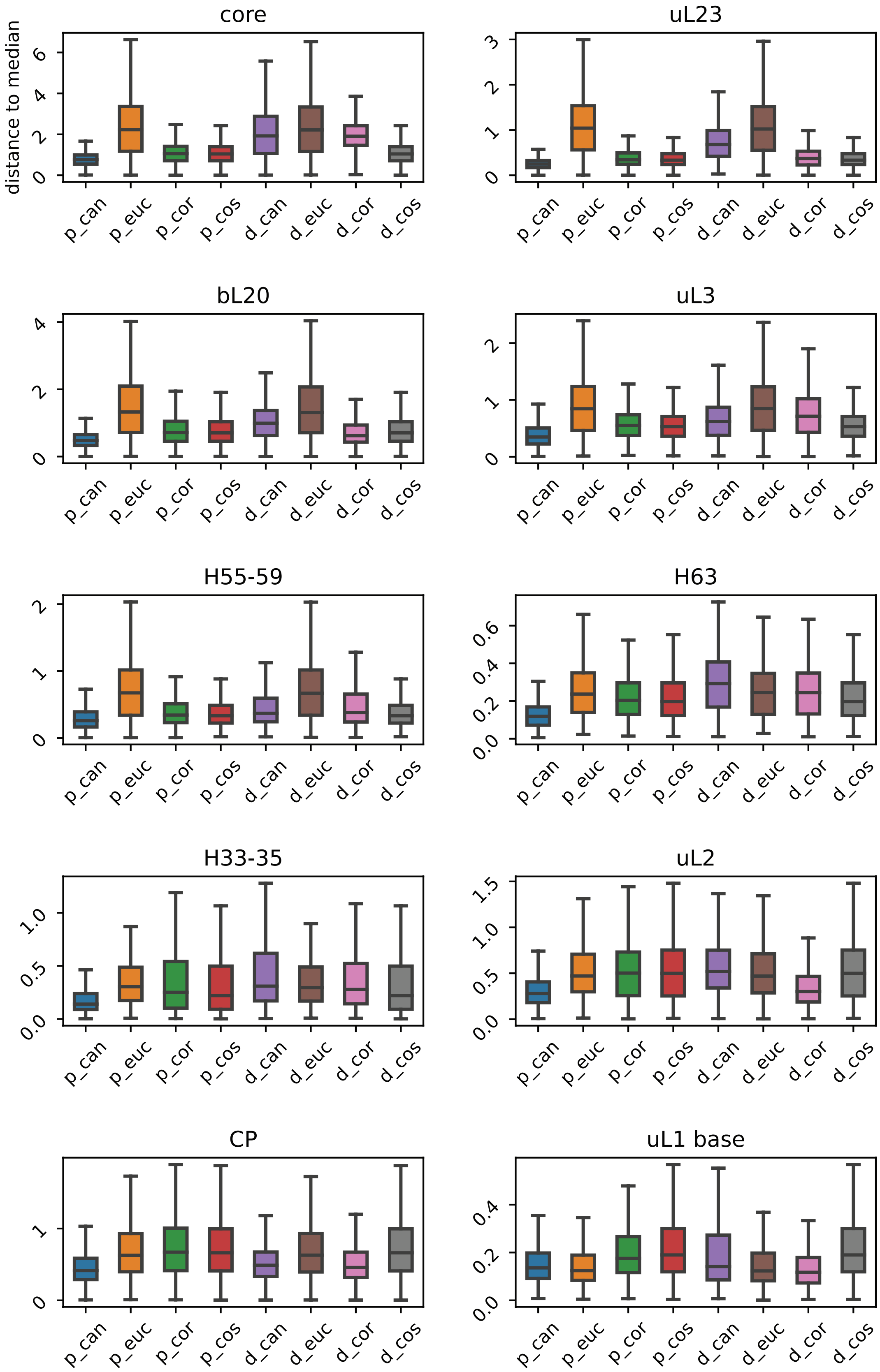
core
uL23
bL20
uL3
H55-59
H63
H33-35
uL2
CP
uL1 base
distance to median
p_can
p_euc
p_cor
p_cos
d_can
d_euc
d_cor
d_cos

**Figure S7 | Boxplot of distance to block median for the 10 blocks from different approaches**
Each boxplot displays the distribution of mean distance to the block median, for each of of 10 blocks, labeled as p- for PCA-UMAP, d- for direct UMAP. can for Canberra, euc for Euclidean, cos for cosine, and cor for correlation.

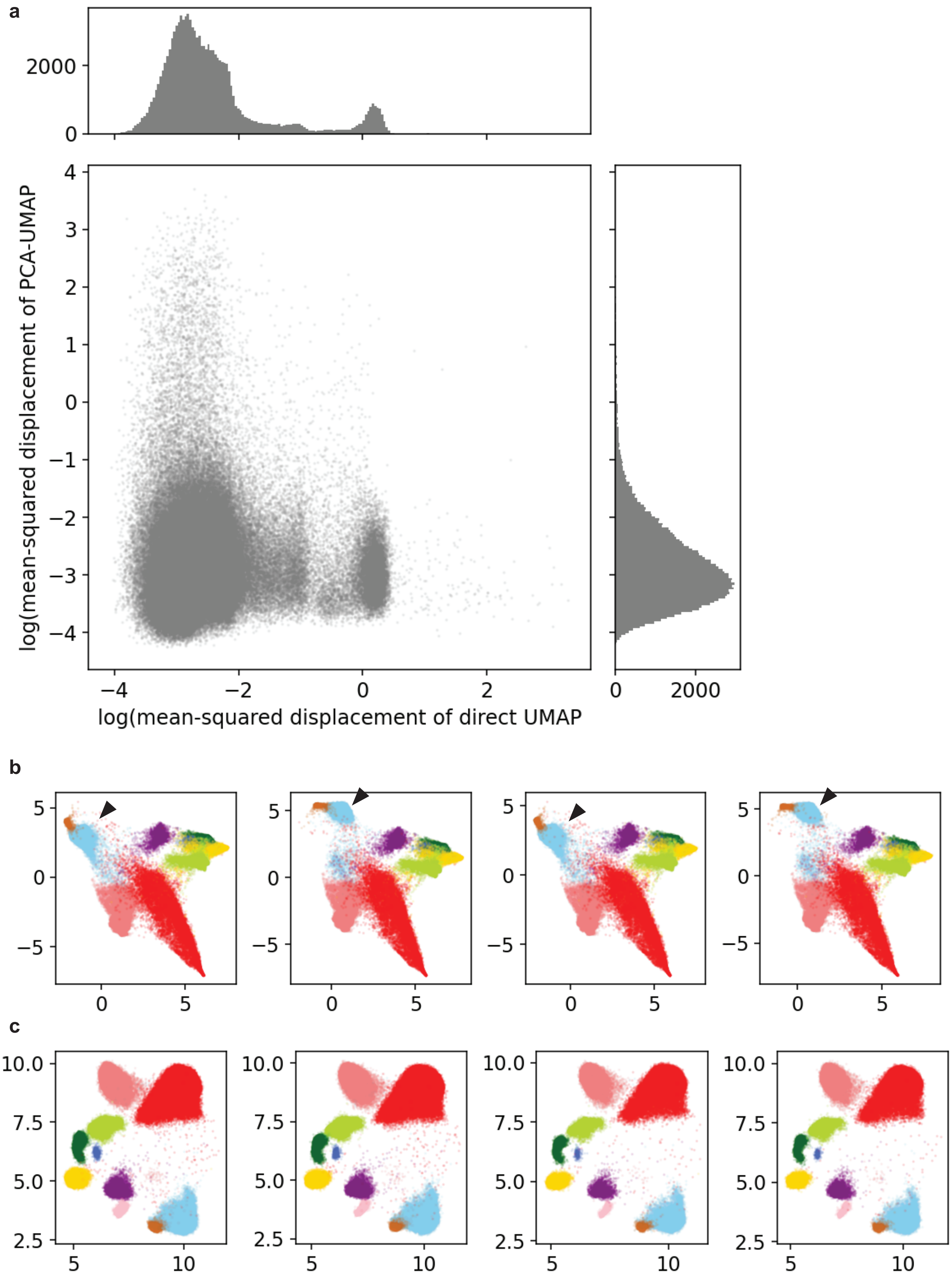
a
b
c
2000
0
4
3
2
1
0
−1
−2
−3
−4
−4
−2
0
2
0
2000
log(mean-squared displacement of PCA-UMAP
log(mean-squared displacement of direct UMAP
5
0
−5
0
5
10.0
7.5
5.0
2.5
5
10

**Figure S8 | Stability test of the UMAP and PCA-UMAP analysis. a**) Mean square displacement of each voxel in 100 independent trials of the direct UMAP (x-axis) and PCA-UMAP (y-axis) calculations, show as a scatterplot with histograms of the marginal distributions at top and right. Representative plots of four consecutive calculations with direct UMAP (**b**) and PCA-UMAP (**c**). The black triangle points out the reorganization of the CP and uL1 blocks in different runs with the direct UMAP method.

**a**

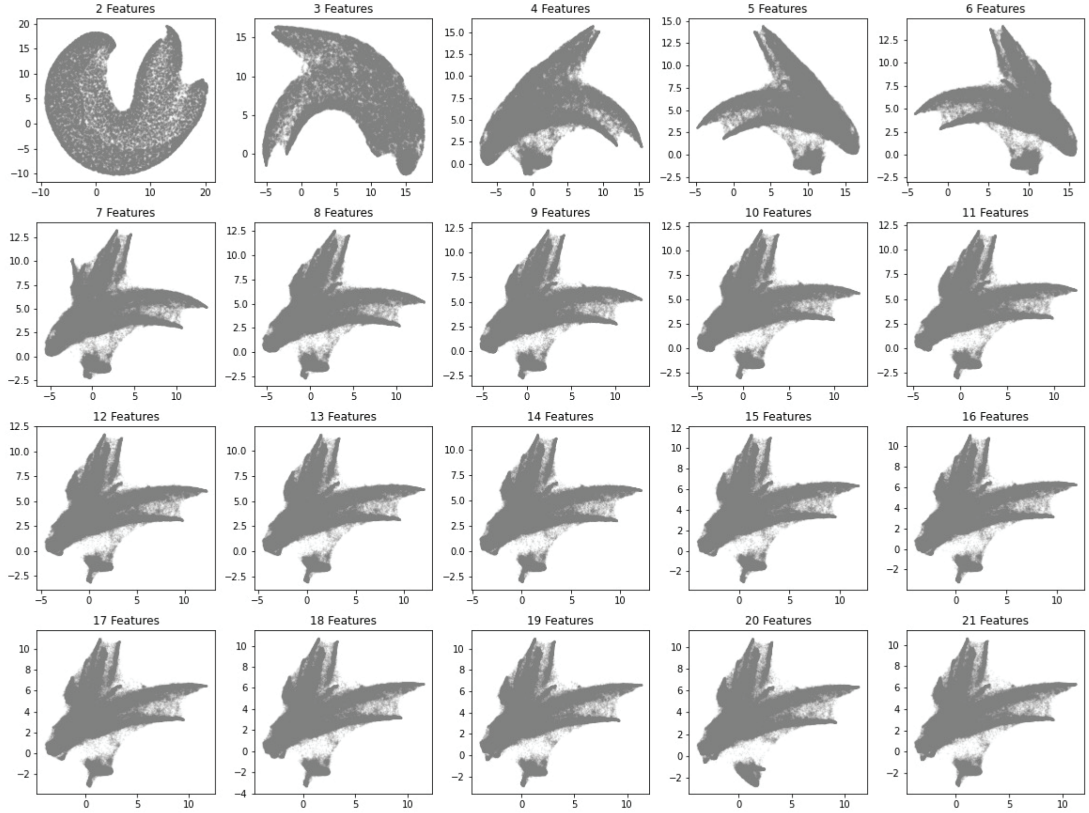


**b**

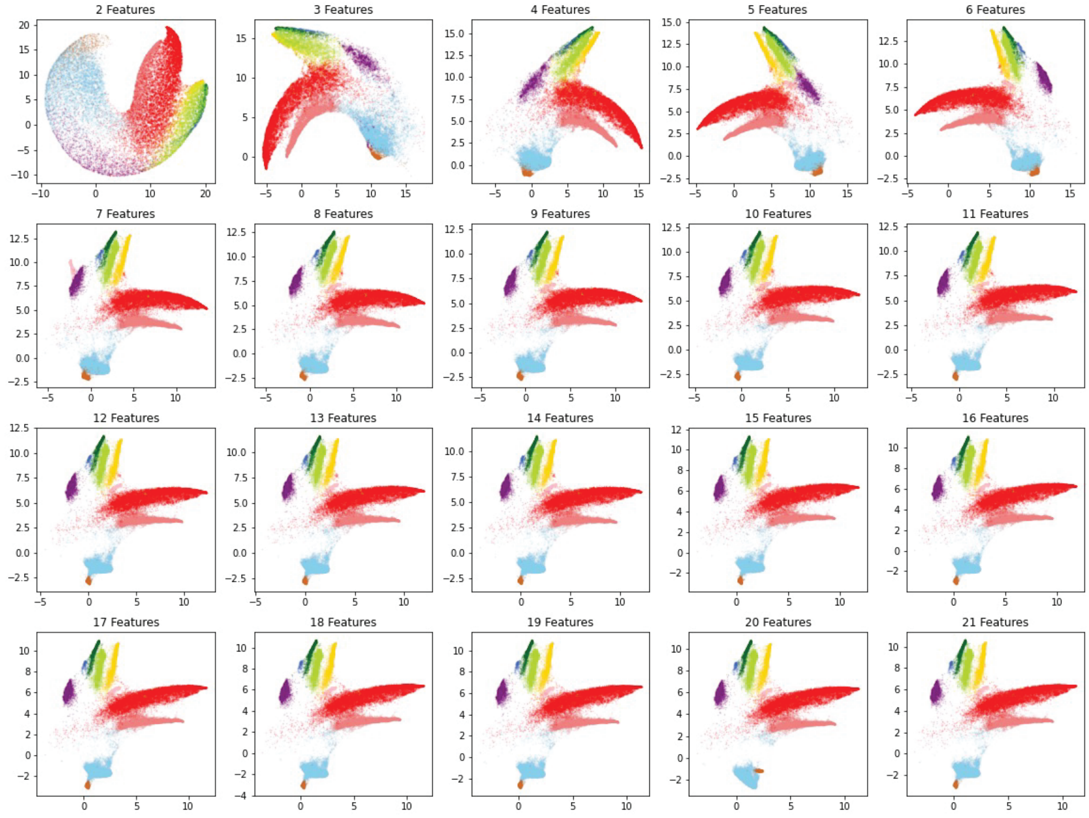


**Figure S9 | PCA-UMAP analysis with different numbers of PCs**

Different numbers of (from 2 to 21) PCs from ΔdeaD dataset were input into UMAP analysis using the Euclidean metric. In the upper panel all voxels as shown in gray, and in the lower panel, the voxels are colored according to Figure 2 for the 10 blocks as a reference.

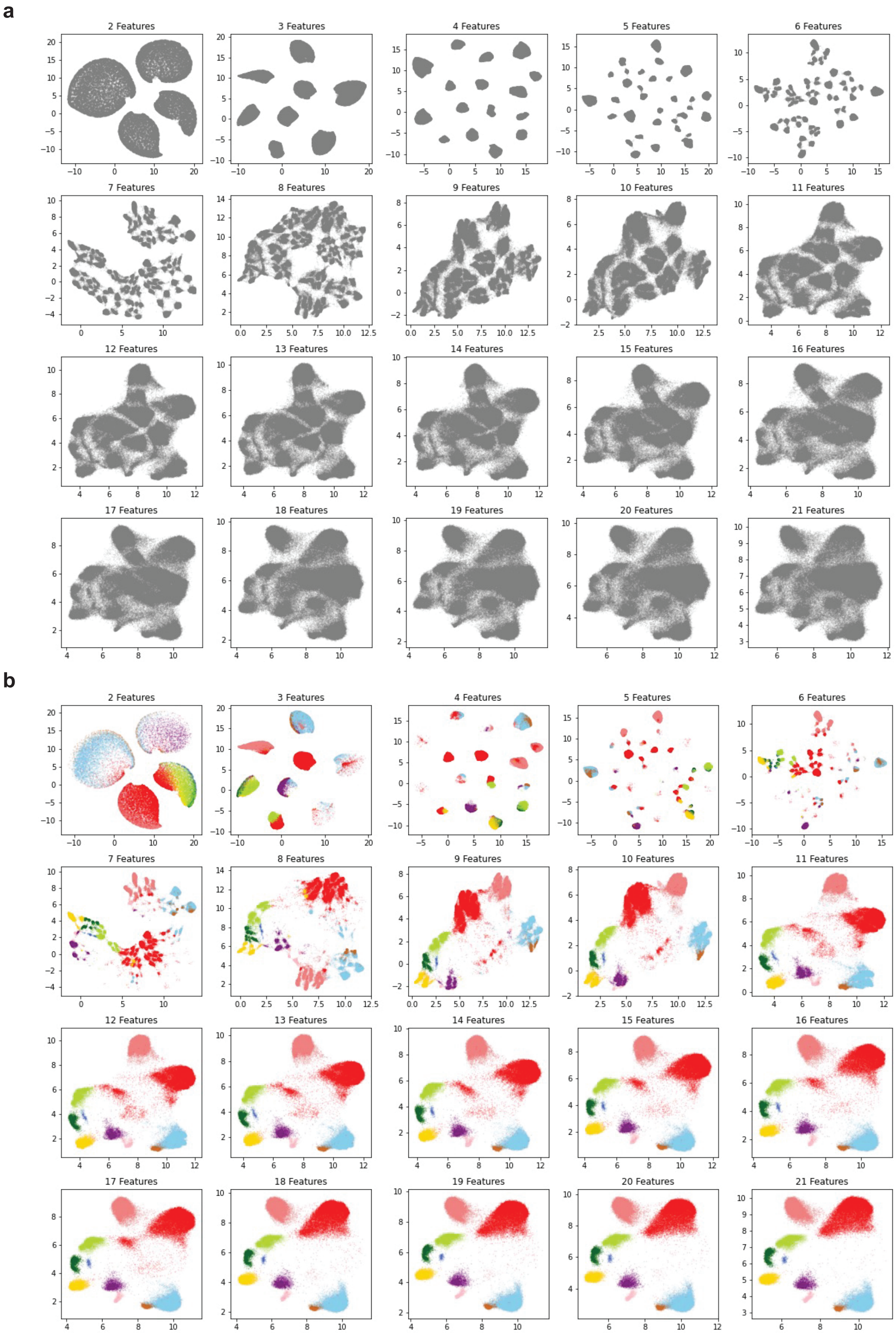


**Figure S10 | PCA-UMAP analysis with different numbers of PCs**

Different numbers of (from 2 to 21) PCs from ΔdeaD dataset were input into UMAP analysis using the Canberra metric.

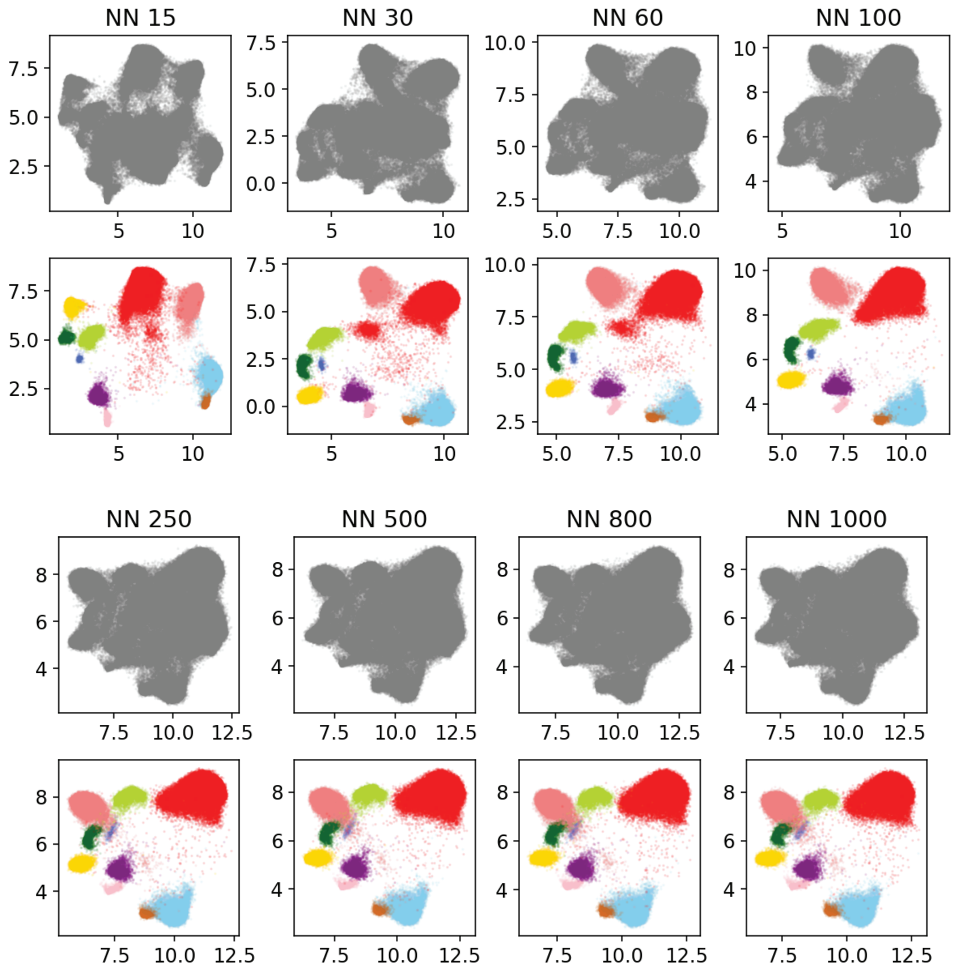


**Figure S11 | PCA-UMAP analysis with different numbers of nearest neighbors**
PCA-UMAP analyisis of the ΔdeaD dataset using the Canberra metric, with different numbers of nearest neighbors for the UMAP (15, 30, 60, 100, 250, 500, 800 and 1000).
In the upper panel all voxels as shown in gray, and in the lower panel, the voxels are colored according to Figure 2 for the 10 blocks as a reference.

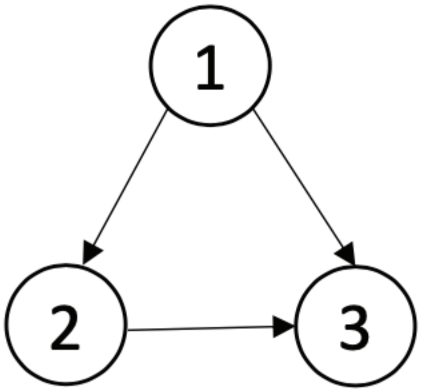


Before pruning

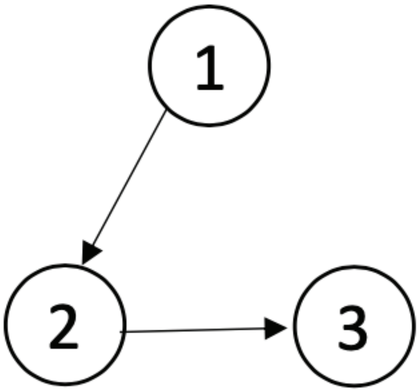


After pruning

**Figure S12 | Scheme for dependency map pruning.** Dependencies that emerge from the quadrant analysis can be represented as directed graphs. To simplify the graph network, edges that short-circuit other paths are pruned, as illustrated for a simple set of 3 nodes with dependences 1->2, 1->3, 2->3.

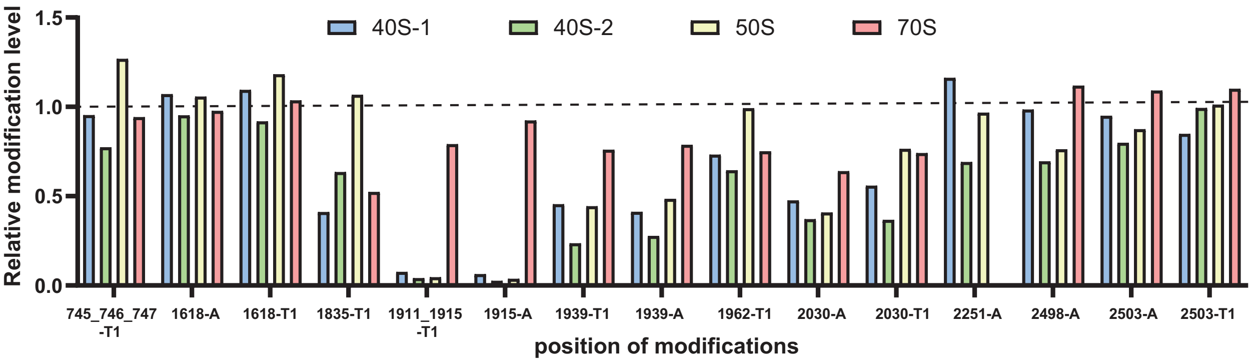


**Figure S13 | Quantification of rRNA modification for ΔdeaD intermediate fractions**

Relative rRNA modification level was quantified by RNA mass spectrometry and displayed as bar graph, grouped by residues and the RNase used for digestion. The bar plot includes two 40S, one 50S and one 70S fraction from the ΔdeaD dataset.
